# Large-scale composite hypothesis testing for omics data analyses

**DOI:** 10.1101/2024.03.17.585412

**Authors:** De Walsche Annaïg, Gauthier Franck, Boissot Nathalie, Charcosset Alain, Mary-Huard Tristan

## Abstract

Composite hypothesis testing using summary statistics is a well-established approach for assessing the effect of a single marker or gene across multiple traits or omics levels. Numerous procedures have been developed for this task and have been successfully applied to identify complex patterns of association between traits, conditions, or phenotypes. However, existing methods often struggle with scalability in large datasets or fail to account for dependencies between traits or omics levels, limiting their ability to control false positives effectively. To overcome these challenges, we present the qch_copula approach, which integrates mixture models with a copula function to capture dependencies between traits or omics, and provides rigorously defined *p*-values for any composite hypothesis. Through a comprehensive benchmark against eight state-of-the-art methods, we demonstrate that qch_copula controls Type I error rates effectively while enhancing the detection of joint association patterns. Compared to other mixture model-based approaches, our method notably reduces memory usage during the EM algorithm, allowing the analysis of up to 20 traits and 10^5^ − 10^6^ markers. The effectiveness of qch_copula is further validated through two application cases in human and plant genetics. The method is available in the R package qch, accessible on CRAN.

## Introduction

Consider a study where the goal is to assess the joint effect of a specific drug treatment in two different tissues. One aims at defining a test procedure that rejects hypothesis *H*_0_ «the drug has no joint effect», when both hypotheses 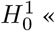 the drug has no effect on tissue 1» and 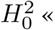 the drug has no effect on tissue 2» are false. This corresponds to a particular case of composite hypothesis testing where the composite *H*_0_ hypothesis to be tested is 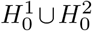. A popular strategy to perform Composite Hypothesis Testing (CHT) is to combine the test statistics and/or *p*-values derived for each of the marginal hypotheses 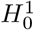 and 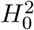 into a single summary statistics. While the *p*-values related to 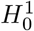 and 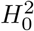 can be obtained using common statistical procedures, providing a suitable summary statistic along with a valid rejection rule (that ensures the control of the false positive rate at the required nominal level) for the test of the composite *H*_0_ hypothesis is not straightforward. The particular CHT problem of the form 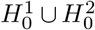 was addressed as soon as the early 80s with the seminal work of (1), and its generalization to the case of testing 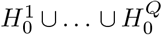 with *Q* ≥ 2 has been investigated by (2).

In the context of omics studies CHT has rapidly become a standard procedure. In human genetics, it has been successfully used for e.g. mediation analysis (3–7), or for the detection of genomic regions that are associated with two diseases such as Prostate Cancer and Type 2 Diabete (8). In these omics applications CHT is typically performed at the gene or marker level, resulting in a large number of simultaneously tested composite hypotheses. A first consequence is the need for CHT methods to explicitly provide *p*-values in order to be combined with multiple testing correction procedures. A second consequence is the opportunity to exploit the large amount of available data to model and infer the relationship between the 2 series of *p*-values, and to account for this relationship in the CHT procedure to better control Type I error rate. More recently, CHT has also been used in the context of integrative genomics to jointly analyse multiple omics traits, such as DNA methylation, copy number variation and gene expression (9, 10), or to the analysis of gene expression kinetics for the detection of differential effects between treatments at two or more consecutive time points (11). When applied to such studies, CHT typically requires the analysis of *Q >* 2 series of *p*-values (i.e. one serie per omics dataset) which led to the development of dedicated procedures.

Most existing methods rely on a similar model where the vector of the *Q* (possibly transformed) *p*-values associated to each gene/marker is assumed to be distributed as a multivariate mixture where each of the 2^*Q*^ components corresponds to a specific combination of 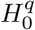 and 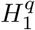 states (7, 9–11).

While versatile, the mixture approach suffers from several limitations. First, as the number of components of the considered mixture model grows exponentially with *Q*, the number of series of *p*-values that can be handled by such CHT procedures is usually restricted to *Q* ≤ 10 (9, 11) or less (e.g. *Q* = 2 and *Q* = 3 in (10) and (7), respectively). Second, while the type I error rate can be controlled through local or multi-class FDR in a mixture model framework (12–14), mixture model-based approaches do not provide *p*-values. This limitation hampers the application of most multiple testing procedures, restricts the use of diagnostic tools such as QQ-plots or histograms, and impedes the ability to compare with alternative methods. Lastly, only a few methods can efficiently handle both a number of *p*-value series *Q* greater than 2 and correlations between series, which may arise, for example, from using the same collection of samples to acquire the different omics measurements.

Based on the same mixture model approach, we propose a new method for CHT called qch_copula that explicitly accounts for the dependence structure across *p*-value series through a copula function (15). The new procedure comes with several important features. First we show how rigorously defined *p*-values can be defined from the mixture model approach and show the connection between adaptive Benjamini-Hochberg FDR control (16) applied to these *p*-values and a local FDR control (12, 13) directly applied to the posteriors obtained from the mixture model. Second, we provide a new implementation of the EM algorithm (17) that significantly alleviate the memory burden of the inference procedure of the approach, extending its application to series of *p*-values as large as *n* = 10^5^*/*10^6^ genes/markers and *Q* = 20. Third, we present the first extensive benchmark study of CHT procedures, comparing 8 recently developed procedures (5, 6, 8–11, 18). While half of the methods do not correctly control for Type I error rate when the *p*-value series are correlated, the qch_copula approach provides accurate Type I error rate control and yields excellent performance in terms of detection power. The procedure is then demonstrated through two application cases: the first in human genetics, where 14 association studies are jointly analyzed to identify new pleiotropic regions associated with psychiatric disorders, and the second in plant genetics, for detecting hotspot regions linked to resistance to multiple viruses in cucumber.

## Materials and methods

This section introduces the framework of composite hypothesis testing then our new method for CHT based on a mixture model approach (Section Model). Inference and the testing procedure are presented in Section Inference and Section Testing composite hypothesis, respectively. The derivation of a memory-efficient EM algorithm is provided in Section Memory-Efficient EM algorithm. The simulation framework used to evaluate the performance of our procedure and its competitors are detailed in Section Simulation framework and Section Comparison with alternatives methods, respectively.

### Composite hypothesis

Assume a collection of *n* items (e.g. genes or SNP) have been tested for their effects in *Q* conditions (e.g. traits, tissues or environments).

We denote by 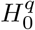 (resp.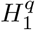) the null (resp. alternative) hypothesis corresponding to test *q* (1 ≤ *q* ≤ *Q*) and consider the set 𝒞:= {0, 1} ^*Q*^ of all possible combinations of null and alternative hypotheses among *Q*. For a given configuration *c* := (*c*_1_, …, *c*_*Q*_) ∈ 𝒞, the joint hypothesis ℋ^*c*^ is defined as

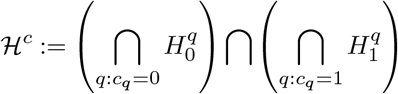

Considering two complementary subsets 𝒞_0_ and 𝒞_1_ satisfying 𝒞_0_ ∪𝒞_1_ = 𝒞 and 𝒞_0_ ∩𝒞_1_ = ∅, we define the composite null and alternative hypotheses ℋ_0_ and ℋ_1_ as

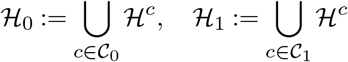

We aim at testing ℋ_0_ versus ℋ_1_ for each of the *n* items.

As an illustration, consider a molecule screening trial where each molecule is tested for 2 desired effects and 2 adverse side effects, referred to as 2 “positive” and 2 “negative” effects respectively in what follows. A molecule is of interest to the experimenter if it has at least one positive effect and no more than one negative effect. Here *Q* = 4, and each configuration has the form *c* = (*c*_1_, …, *c*_4_) where the first 2 elements *c*_1_, *c*_2_ correspond to the 2 tests for positive effects. The configurations of interest for the experimenter are

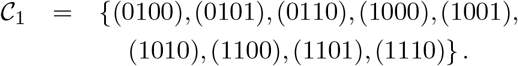

The complementary set is

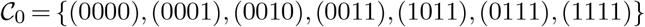

corresponding to configurations where either the molecule has no positive effect (first 4 configurations) and/or 2 negative effects (last 4 configurations). Testing composite hypothesis ℋ_0_ boils down to testing whether the (unknown) configuration of the molecule under study belongs to 𝒞_0_ or not. We stress out that in this example, the alternative composite hypothesis ℋ_1_ actually requires that at least one of the two null hypotheses of negative effects be true, exemplifying how complex combinations of basic hypotheses may be tested through the proposed setting.

### Model

Let 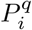 denote the *p*-value obtained for test *q* on item *i*, and let 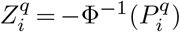 represent its negative probit transform, where Φ stands for the standard Gaussian cumulative distribution function (cdf). The vector 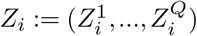 is referred to as the z-score profile of item *i*.

Each item *i* is associated with a latent vector 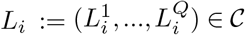, where 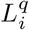 is the binary variable indicating whether the null hypothesis 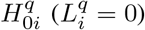 or its alternative hypothesis 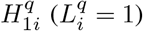 holds. As such *L*_*i*_ corresponds to the unobserved label of item *i*, i.e the true configuration to which *i* belongs to. Under the assumption of independence between items, the z-score profile follows a mixture model with 2^*Q*^ components, i.e. one for each configuration, and can be expressed as:

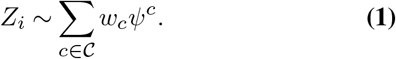

where *ψ*^*c*^ is the distribution of *Z*_*i*_ conditional on *L*_*i*_ = *c*, and the 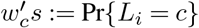 represent the mixing proportions.

Each component *ψ*^*c*^ corresponds to a multivariate distribution over **R**^*Q*^ that can be written in terms of univariate marginal distribution functions and a so-called copula function that describes the dependence structure between the *Q* z-scores (15):

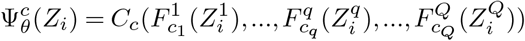

where 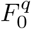 (resp. 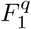) is the marginal cdf of 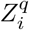 conditional on 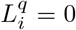 (resp.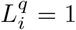). In what follows, we will assume that the copula function is common to all components and depends on a finite set of unknown parameters *θ*, that is, *C*_*c*_ = *C*_*θ*_ ∀ _*c*_ ∈ 𝒞. The corresponding density function for component *c* is then given by:

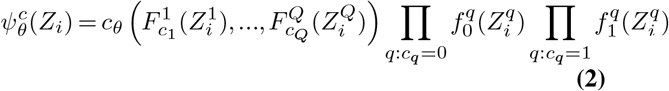

where 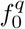 (resp.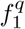) is the marginal density of 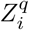 conditional on 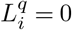 (resp.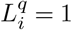). Since z-scores are obtained from *p*-values whose distribution is known to be uniform over [0, 1] under *H*_0_, all distributions are known: one has 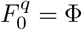 for all *q*.

The expression obtained in Eq. (2) provides some hints about the complexity of the inference task: estimating the 2^*Q*^ conditional distributions *ψ*_*c*_ of the mixture model Eq. (1) actually reduces to determining the *Q* univariate cumulative distributions 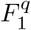 and the copula parameter *θ*. In practice, one needs to choose a specific form of the copula distribution. Here we considered Gaussian copula due to its flexibility in specifying distinct correlation levels between each pair of variables. The density function of the Gaussian copula is given by:

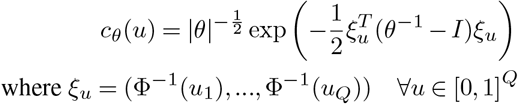

and where *θ* is the (*Q × Q*) correlation matrix associated with the Gaussian copula.

Note that the scenario in which *θ* varies across components, and where the alternative distributions 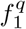 are modeled as a Gaussian distribution, aligns with the methodology proposed by (10), which employs a mixture model of multivariate Gaussian distributions. Conversely, the specific case where *θ* = *I*_*Q*_ and where the alternative distributions 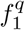 are inferred in a non-parametric way, corresponds to the independent mixture model introduced in (11).

### Inference

The unknown parameters of model Eq. (1) are the distributions 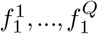, the copula parameter *θ* and the proportions *w*_*c*_. Following (9) and (11), the inference procedure can be split into two steps:

1. Get an estimate of marginal densities 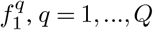;
2. Substitute the estimates 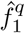 into Equation Eq. (1) and estimate both the proportions *w*_*c*_ and the copula parameter *θ* using maximum likelihood estimation.

#### .1. Step 1: Inference of marginal distributions

Combining Model Eq. (1) with the definition of the *ψ*_*c*_ Eq. (2), one has

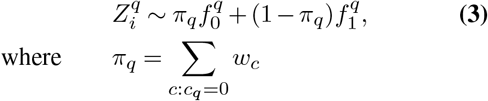

that is the marginal distribution of 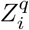 is also a mixture model. Since 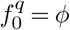 for all *q*’s, with *ϕ* the standard Gaussian density function, one needs to estimate 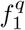 and *π*_*q*_ only.

The null proportions can be directly derived by applying the following estimator:

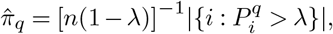

where *λ* ∈ [0, 1] is a tuning parameter that can be determined through bootstrap (19). The estimated proportion 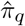 can then be plugged into equation Eq. (3), and the alternative distribution 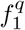 can then be inferred through a nonparametric procedure using a kernel method (13). In terms of computational burden, this procedure can be executed for each of the *Q* components in parallel.

#### Step 2: Inference of the configuration proportions and the copula parameter

We now turn to the problem of inferring the copula parameter *θ* and the weights *w*_*c*_ using maximum likelihood estimation. Once the kernel estimates 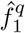 are substituted in mixture model Eq. (1), the inference of the remaining parameters can be efficiently performed using a standard EM algorithm (17) using the full set of items. In the present case, it is possible to obtain explicit update equations for both *θ* and the *w*_*c*_’s. Denoting 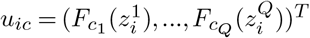 and 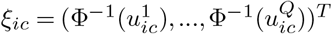, one has :

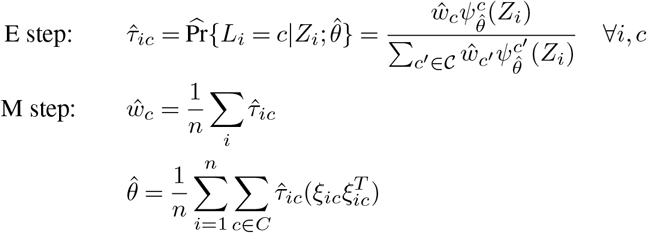

See Section 1 of the Supplementary Material for a detailed explanation of these derivations. Note that the expression of matrix *θ* ensures its positiveness. If applied directly, the (E) step involves computing and storing a posterior matrix of size *n ×* 2^*Q*^. Although computational effort cannot be avoided, we demonstrate in Section that memory storage can be considerably reduced without loss of information.

### Testing composite hypothesis

So far, methods relying on mixture model Eq. (1) (9–11) did not result in strictly valid test procedures as they did not provide a *p*-value. We present here how well-defined *p*-values can be derived in this framework.

Let *c* be the (unknown) configuration of the item under consideration, and 𝒞_0_ and 𝒞_1_ be any two complementary subsets of 𝒞, and consider testing

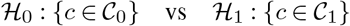

The mixture model Eq. (1) can be written in the following form:

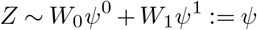

where 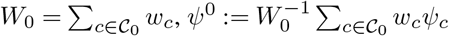 (resp. for *W*_1_ and *ψ*^1^). We consider the posterior

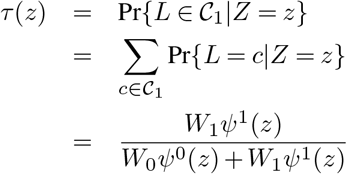

as a test statistic. The corresponding *p*-value is then:

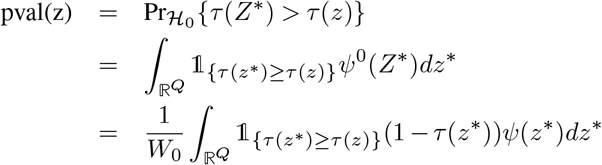

where the last equality comes from the definition of *τ*. In practice, estimates of posteriors

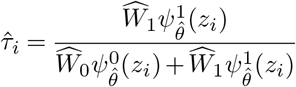

can be computed for each item *i*. Using these estimates and approximating the integral by its empirical counterpart, one gets:

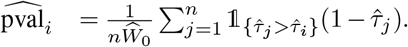

Having access to *p*-values provides several advantages over existing methods that rely on posteriors only, from combining the testing procedure with any correction method for multiple testing to checking the *p*-value distribution for quality control and providing graphical displays such as Volcano or Manhattan plots.

Beyond the empirical expression provided above, a theoretical equivalence can be established between *p*-values 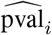 corrected for multiple testing using the adaptive Benjamini Hochberg procedure (16) on one side, and the estimation of lFDR from the posteriors as presented in (12) and (13) on the other side. This equivalence is demonstrated in the Section 2 of the Supplementary Material.

### Memory-Efficient EM algorithm

The classical EM algorithm (17) implementation requires the computation and the storage of the matrix *T* = (*τ*_*ic*_) in (E) step, which becomes cumbersome whenever *n* and/or *Q* are large: assuming e.g. *n* = 10^5^ and *Q* = 15, the matrix *T* requires 26 GB of storage. This storage may be reduced by analyzing how these posteriors are used in the (M) step. For instance, the update of *w*_*c*_ at iteration (*t* + 1) can be reformulated as:

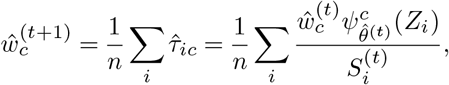

where 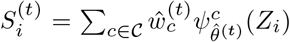. Our reduced memory burden implementation of the EM algorithm works as follows: in the (E) step only the quantities 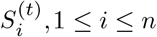 are computed and saved; then the 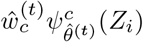 are calculated on the fly for updating the weights 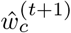 in the (M) step. A same procedure can be derived for the copula parameter 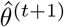 update, see Suppl. Mat. Section 3 for details.

This version of the EM algorithm retains the exact formulas of the original algorithm while significantly reducing its memory footprint from *O*(*n ×* 2^*Q*^) to *O*(*n* + 2^*Q*^). In our previous example (*n* = 10^5^, *Q* = 15) the memory storage is downsized from 26 GB to 1 MB. This optimization makes it possible to apply our procedure to cases where *n* ranges from 10^5^ to 10^6^ and with up to *Q* = 20 conditions.

In what follows we will refer to the methodology presented so far (including mixture models and copulas, the use of an efficient EM algorithm for the inference and the derived *p*-values for the CHT procedure) as the qch_copula approach.

### Simulation framework

We evaluated and compared the performance of our qch_copula method with existing methods in various simulated scenarios. Hereafter, we will refer to items and conditions as “markers” and “traits”, mimicking a multi-trait genome-wide association study. Each simulated dataset consists of a *n × Q* matrix of z-scores generated as follows. Markers are clustered into *K* + 1 clusters. Each cluster is characterized by a configuration *c*, i.e. all markers of cluster *k* share the same configuration *c*_*k*_ = (*c*_*k*1_, …, *c*_*kQ*_). One has

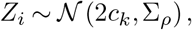

where Σ_*ρ*_ is a *Q × Q* correlation matrix where all covariances between traits are set to *ρ*. Note that *µ*_*iq*_ = 0 if 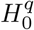 holds for item *i*, and *µ*_*iq*_ = 2 otherwise. Assuming that matrix Σ_*ρ*_ is common to all clusters, a given cluster *k* is characterized by its associated number of markers and associated configuration vector. Figure 1 provides an illustrative example of the z-score matrix with items grouped into the *K* + 1 clusters, each characterized by a configuration.

**Fig. 1.**
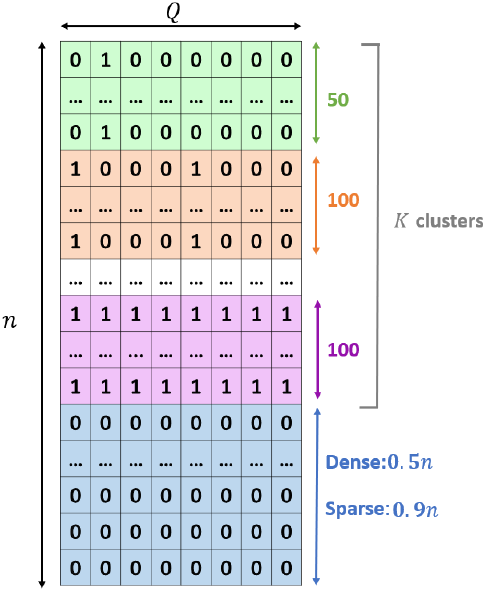
Schematic illustration of the *n × Q* z-score matrix used in the simulation study. Each row represents an item (e.g. a marker), and each column corresponds to one of the *Q* conditions (e.g. a trait). Items are grouped into clusters, each corresponding to a configuration of associations across conditions. The last cluster, represented in blue, corresponds to items for which the complete null configuration is true.

The aim of the statistical analysis performed on these simulated datasets is then to identify markers that are associated with at least *q* traits (with different values of *q* to be considered). Note that we are not interested in inferring the true configuration of the markers.

We considered different values for *Q* (2,8,16), four levels of correlation (*ρ* = 0, 0.3, 0.5, 0.7), and two scenarios: *sparse* or *dense*. In the *sparse* scenario, a large proportion (i.e. ≥ 0.9) of the markers were associated with none of the traits, whereas in the dense case, half of the markers were associated with at least one trait. For each setting (i.e., a combination of a number of traits, a correlation level and a scenario), simulations were repeated 25 times. In total, 3 *×* 4 *×* 2 = 24 settings were investigated (see Table 8 in Suppl. Mat. For details). All simulations were performed using *n* = 10^5^.

We provide the details for generating one dataset in Setting 1: *Q* = 2, *ρ* = 0, *sparse*, see Suppl. Mat. Section 4 for the description of other simulation settings. The dataset includes clusters of markers associated with different configurations, as summarized in Table 1. Specifically, twenty clusters of 50 markers associated with a single randomly chosen trait were simulated. Similarly, twenty clusters of 50 markers associated with the two traits were simulated. The process was repeated to generate ten clusters of 100 markers associated with one trait and ten clusters of 100 markers associated with two traits. A last cluster containing all remaining markers corresponding to the complete null configuration, i.e. *c* = (0, 0), was generated. This last cluster included 96,000 markers, i.e., 96% of the total number of markers.

**Table 1.** Summary of cluster composition for Setting 1 (*Q* = 2, *ρ* = 0, sparse scenario).

| Number of clusters | Cluster size | Configuration $c$ |
| --- | --- | --- |
| 20 | 50 | (1, 0) or (0, 1) |
| 10 | 100 | (1, 0) or (0, 1) |
| 20 | 50 | (1, 1) |
| 10 | 100 | (1, 1) |
| 1 | 96,000 | (0, 0) |

To assess the robustness of the proposed methods in situations involving non-independent items, we implemented an additional simulation framework incorporating within trait dependence among markers. Specifically, markers were grouped into blocks such that p-values within each block were correlated, mimicking the local dependence structure typically observed in e.g. genome-wide association studies due to linkage disequilibrium. We considered a spatial correlation level of *ξ* = 0.3 within blocks, combined with the same 24 simulation settings described above. A detailed description of this simulation setting is provided in Supplementary Material Section 4.

### Comparison with alternative methods

Our benchmarking study evaluated eight recently proposed methods: DACT (5), HDMT (6), PLACO (8), adaFilter (18), IMIX (10), c-csmGmm (7), Primo (9) and qch (11). The first three methods-DACT, HDMT, and PLACO - use test statistics that combine two *p*-values per item and are restricted to scenarios involving no more than two test series. In contrast, IMIX, c-csmGMM, Primo and qch rely on mixture models similar to the one described in Equation Eq. (1) and can theoretically

handle cases with more than two test series. However, in practice, both IMIX and c-csmGmm had prohibitive computational costs and/or the associated R package did not handle cases where *Q >* 3 and *Q >* 2,respectively. The adaFilter method takes a different approach, testing a specific composite null hypothesis of the form “fewer than *q* hypotheses among the *Q* tested are non-null”, using an adaptive filtering multiple testing procedure with no restriction on *Q*. Among all considered methods, only PLACO, IMIX, Primo, c-csmGmm and qch_copula explicitly account for dependencies between *p*-values series. As Primo only provides an estimate of the local FDR for each marker, we relied on the equivalence between local FDR (lFDR) estimation and *p*-values based on posteriors proven in Section to derive *p*-values for each marker for this method. Additionally, the data simulation process described in the previous section corresponds to the model described by Equations Eq. (1) and Eq. (2) with the marginal alternative distributions 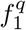 being the Gaussian distribution 𝒩 (2, 1) and the copula parameter *θ* being Σ_*ρ*_. This setup may be beneficial in terms of detection power to IMIX, Primo and qch_copula which rely on the same mixture model. However, all benchmarked methods should ensure a control of Type I error rate at the nominal level.

## Results

### Simulations

Our evaluation began by assessing the ability of the different methods to control Type I error (T1E) at the specified significance level and to detect true significant markers for *Q* = 2. We then extended the comparison to *Q* = 8 and *Q* = 16 for methods capable of handling larger cases.

Additional information regarding the accuracy of qch_copula to estimate the proportions of items under the null hypothesis in each condition are provided in Section 6 of the Supplementary Materials.

#### Type I error rate control

In settings 1-8, *n* = 10^5^ markers were tested for *Q* = 2 traits, with the objective of identifying markers associated with both traits. FDR control at a nominal level of 5% was applied using the adaptive BH procedure to generate the final list of candidate markers for each method. The results for the case *ρ* = 0 are presented in Table 10 in Supp. Mat. Most methods successfully control the FDR at the nominal level, with minor deviations observed for PLACO and c-csmGmm, which slightly exceed the target (approximately 0.08). In contrast, DACT-JC displays a substantial inflation of the FDR, reaching values around 0.25.

The results for the case *ρ* = 0.3 and *ρ* = 0.5 are summarized in Table 2 and Table 11 in Supp. Mat., revealing significant departures from the nominal Type I error rate for most methods.

**Table 2.** False discovery rate of the procedures for the sparse and dense scenarios with *Q* = 2 and *ρ* = 0.3 (settings 1 and 2).

| Method | Sparse | Dense |
| --- | --- | --- |
| DACT_Efron | 0 (0) | 0 (0) |
| DACT_JC | 0 (0) | **0.08 (0.017)** |
| DACT_NO | 0.032 (0.03) | 0.014 (0.015) |
| HDMT | **0.255 (0.018)** | **0.129 (0.008)** |
| PLACO | **0.148 (0.091)** | 0.058 (0.027) |
| adaFilter | **0.1 (0.015)** | **0.087 (0.008)** |
| qch | **0.256 (0.018)** | **0.145 (0.008)** |
| IMIX | **0.595 (0.128)** | **0.222 (0.094)** |
| c-csmGmm | **0.186 (0.085)** | **0.067 (0.019)** |
| Primo | 0.046 (0.021) | **0.079 (0.01)** |
| qch_copula | 0.045 (0.021) | 0.057 (0.012) |

Note: For each procedure the FDR (averaged over 25 runs) are displayed. Numbers in brackets correspond to standard errors. False discovery rate above 0.05 are in boldface.

As expected the qch_copula method exhibited substantial improvements over the original qch approach (which assumes conditional independence between *p*-value series), highlighting the importance of accounting for the dependency structures. More generally, only DACT_Efron and qch_copula consistently achieved proper FDR control, maintaining estimated FDR values close to 0.05 across all scenarios (settings 3-6:*ρ* = 0.3 and *ρ* = 0.5).

Similar results were observed when the correlation between traits was higher (*ρ* = 0.7; see Table 12 in Supp. Mat.) and in the presence of dependence structure between items (*ξ* = 0.3; see Tables 13-16 in Supp. Mat.), with increased FDR inflation among methods that failed to adequately control the FDR.

We further investigated TIE control in settings 9-16 and 17-24, corresponding to cases with *Q* = 8 and *Q* = 16, respectively, to identify markers associated with at least 2, 4, 8, 14 or 16 traits. Only Primo, adaFilter and qch_copula were included in these analyses, as the other methods are restricted to scenarios where *Q* = 2 or *Q* ≤ 3. The results for *ρ* = 0.3 are presented in Table 3, while additional results for *ρ* = 0, 0.5, and 0.7 are provided in Tables 17–19 of the Suppl. Mat. In settings 9–16 (*Q* = 8), all three methods demonstrated effective T1E control when the correlation between traits was low to moderate (i.e., *ρ* = 0 and *ρ* = 0.3), with Primo and adaFilter generally exhibiting conservative behavior relative to the nominal level. At higher correlation levels (*ρ* = 0.5 and *ρ* = 0.7), small to moderate deviations from the nominal T1E were observed. For qch_copula, T1E remained below 0.07 in most scenarios, with the highest observed value of 0.116 when testing for association with at least 8 traits. In comparison, Primo and adaFilter showed T1E up to 0.142 and 0.246, respectively.

**Table 3.** Performance of the procedures Primo, adaFilter and qch_copula for the settings 11-12 (*Q* = 8) and 19-20 (*Q* = 16) with *ρ* = 0.3.

| $Q$ | Scenario | Test | Primo | | adaFilter | | qch_copula | |
| --- | --- | --- | --- | --- | --- | --- | --- | --- |
|  |  |  | FDR | Power | FDR | Power | FDR | Power |
| 8 | Dense | at least 2 | 0.008 (0.001) | 0.143 (0.006) | 0.018 (0.001) | 0.317 (0.003) | 0.051 (0.001) | 0.609 (0.003) |
|  |  | at least 4 | 0.005 (0.001) | 0.126 (0.007) | 0.006 (0.001) | 0.148 (0.003) | 0.044 (0.002) | 0.534 (0.006) |
|  |  | at least 8 | 0.048 (0.006) | 0.125 (0.007) | 0.017 (0.007) | 0.033 (0.003) | 0.05 (0.006) | 0.186 (0.01) |
|  | Sparse | at least 2 | 0.012 (0.004) | 0.109 (0.008) | **0.063 (0.005)** | 0.279 (0.008) | 0.054 (0.004) | 0.388 (0.007) |
|  |  | at least 4 | 0.008 (0.004) | 0.108 (0.009) | 0.015 (0.004) | 0.145 (0.005) | 0.053 (0.004) | 0.405 (0.014) |
|  |  | at least 8 | **0.079 (0.019)** | 0.092 (0.009) | 0.026 (0.021) | 0.032 (0.01) | **0.068 (0.023)** | 0.166 (0.019) |
| 16 | Dense | at least 2 | - | - | 0.02 (0.001) | 0.278 (0.002) | 0.052 (0.002) | 0.646 (0.004) |
|  |  | at least 4 | - | - | 0.007 (0.001) | 0.166 (0.002) | 0.057 (0.002) | 0.678 (0.006) |
|  |  | at least 8 | - | - | 0.006 (0.001) | 0.094 (0.004) | 0.057 (0.002) | 0.659 (0.007) |
|  |  | at least 14 | - | - | 0 (0.001) | 0.015 (0.002) | 0.04 (0.004) | 0.548 (0.016) |
|  |  | at least 16 | - | - | 0.036 (0.028) | 0.009 (0.003) | **0.068 (0.013)** | 0.14 (0.02) |
|  | Sparse | at least 2 | - | - | **0.076 (0.005)** | 0.248 (0.005) | **0.072 (0.004)** | 0.528 (0.008) |
|  |  | at least 4 | - | - | 0.025 (0.005) | 0.166 (0.005) | **0.082 (0.004)** | 0.626 (0.01) |
|  |  | at least 8 | - | - | 0.007 (0.006) | 0.097 (0.008) | **0.068 (0.006)** | 0.568 (0.018) |
|  |  | at least 14 | - | - | 0 (0) | 0.016 (0.005) | 0.036 (0.01) | 0.37 (0.027) |
|  |  | at least 16 | - | - | 0.015 (0.031) | 0.012 (0.007) | **0.101 (0.041)** | 0.138 (0.031) |

Note: For each procedure the FDR and the detection power (averaged over 25 runs) are displayed. Numbers in brackets correspond to standard errors. False discovery rate above 0.05 are in boldface.

The results for *ρ* = 0.3 under structured dependence between items are presented in Table 4, with additional results for *ρ* = 0, 0.5, and 0.7 provided in Tables 20–22 of the Suppl. Mat. Dependence between items had no impact on FDR control for adaFilter. In contrast, both Primo and qch_copula exhibited improved FDR control under dependence, particularly in settings where some inflation had been observed in the independent case. In particular, for qch_copula, the FDR decreased significantly in the most challenging case (testing for association with at least 8 traits), from approximately 0.11 under independence to below 0.065 when item dependence was introduced.

**Table 4.** Performance of the procedures Primo, adaFilter and qch_copula for the simulation settings 11s-12s (*Q* = 8) and 19s-20s (*Q* = 16) with *ρ* = 0.3 and *ξ* = 0.3.

| $Q$ | Scenario | Test | Primo | | adaFilter | | qch_copula | |
| --- | --- | --- | --- | --- | --- | --- | --- | --- |
|  |  |  | FDR | Power | FDR | Power | FDR | Power |
| 8 | Dense | at least 2 | 0.006 (0.005) | 0.088 (0.073) | 0.018 (0.002) | 0.316 (0.009) | 0.048 (0.005) | 0.611 (0.016) |
|  |  | at least 4 | 0.003 (0.003) | 0.077 (0.064) | 0.006 (0.002) | 0.146 (0.009) | 0.033 (0.004) | 0.509 (0.022) |
|  |  | at least 8 | 0.031 (0.026) | 0.074 (0.062) | 0.019 (0.008) | 0.031 (0.007) | 0.031 (0.01) | 0.151 (0.025) |
|  | Sparse | at least 2 | 0.008 (0.008) | 0.068 (0.058) | **0.063 (0.006)** | 0.278 (0.022) | 0.055 (0.009) | 0.418 (0.051) |
|  |  | at least 4 | 0.005 (0.004) | 0.068 (0.057) | 0.015 (0.005) | 0.146 (0.014) | 0.049 (0.01) | 0.426 (0.077) |
|  |  | at least 8 | 0.048 (0.05) | 0.053 (0.05) | 0.018 (0.019) | 0.027 (0.012) | 0.056 (0.039) | 0.119 (0.078) |
| 16 | dense | at least 2 | - | - | 0.02 (0.002) | 0.276 (0.011) | 0.054 (0.004) | 0.667 (0.015) |
|  |  | at least 4 | - | - | 0.007 (0.002) | 0.164 (0.009) | 0.053 (0.005) | 0.676 (0.02) |
|  |  | at least 8 | - | - | 0.006 (0.003) | 0.09 (0.009) | 0.05 (0.005) | 0.584 (0.031) |
|  |  | at least 14 | - | - | 0.001 (0.002) | 0.014 (0.004) | 0.019 (0.002) | 0.354 (0.042) |
|  |  | at least 16 | - | - | 0.027 (0.037) | 0.008 (0.005) | 0.042 (0.021) | 0.061 (0.038) |
|  | Sparse | at least 2 | - | - | **0.074 (0.008)** | 0.25 (0.021) | **0.076 (0.009)** | 0.519 (0.031) |
|  |  | at least 4 | - | - | 0.023 (0.005) | 0.168 (0.018) | **0.08 (0.011)** | 0.575 (0.045) |
|  |  | at least 8 | - | - | 0.005 (0.005) | 0.095 (0.025) | **0.071 (0.013)** | 0.42 (0.06) |
|  |  | at least 14 | - | - | 0 (0) | 0.014 (0.009) | 0.02 (0.01) | 0.168 (0.061) |
|  |  | at least 16 | - | - | 0.002 (0.008) | 0.008 (0.007) | 0.056 (0.06) | 0.048 (0.052) |

Note: For each procedure the FDR and the detection power (averaged over 25 runs) are displayed. Numbers in brackets correspond to standard errors. False discovery rate above 0.05 are in boldface.

In settings 17–24 (*Q* = 16), a slight inflation of the Type I error (T1E) was observed for qch_copula when *ρ* = 0.3 under the sparse scenario, although values generally remained below 0.08. At higher correlation levels (*ρ* = 0.5 and *ρ* = 0.7), deviations from the nominal T1E were observed for both adaFilter and qch_copula. Specifically, qch_copula exceeded 0.1 of FDR in 4 out of 20 cases, with a maximum of 0.24, while adaFilter exceeded 0.1 in 8 out of 20 cases (with a maximum of 0.28). The dependency between items resulted in an FDR not exceeding 0.089 for qch_copula in all scenarios, and had no or little impact on adaFilter.

We do not report results for Primo in settings 17-24 as it encountered significant computational challenges, either exhausting available memory or requiring excessively long runtimes, exceeding 24 hours.

#### Detection power

For simulations with *Q* = 2 and focusing on the methods that effectively controlled the FDR across all scenarios, DACT_Efron yielded highly conservative results, with detection power equal to zero. In contrast the qch_copula method achieved detection power ranging from 0.03 to 0.124 depending on the scenario, see Tables 9-16 in Suppl. Mat.

The results of the power detection analysis for settings 11-12, 11s-12s (*Q* = 8) and 19-20, 19s-20s (*Q* = 16) are presented in Table 3 and 4 for the three methods that were able to scale. In both the sparse and dense scenarios the performance of Primo was stable whatever the tested composite hypothesis ℋ_1_, whereas adaFilter showed a marked decrease in power when moving from ℋ_1_: “at least 2” to ℋ_1_: “at least 8” or “at least 16”. This trend is consistent with the conservative behavior observed for adaFilter in the previous section. The qch_copula method exhibited a decline in power when testing the most stringent hypothesis (i.e., ℋ1: “at least *Q*”). However, qch_copula consistently showed a substantial improvement in power over Primo and adaFilter across all settings. For instance, in the sparse scenario with *Q* = 16 and ℋ1: “at least 8”, qch_copula achieved a power that was approximately six times higher than that of adaFilter.

To provide a more comprehensive evaluation of the methods performance, we include Precision–Recall (PR) curves for the case *ρ* = 0.3 (Fig. 4-9 in Suppl. Mat.). These curves complement the FDR and power analyses by illustrating the trade-off between precision (1 – FDR) and recall (power) across a range of decision thresholds. One can observe contrasted behaviors between methods, with PLACO, c-csmGmm, and IMIX exhibiting noticeably more erratic behaviors in sparse scenarios, with greater variability across simulations. All other methods follow similar PR trajectories, but differ in their choice of significance threshold. This pattern suggests that the observed variations in power and FDR are largely driven by thresholding behavior rather than inherent limitations in the methods’ ability to discriminate between alternative and null hypotheses items. This phenomenon is particularly pronounced in the setting with *ρ* = 0 (Fig. 10–13 in Suppl. Mat.).

Similar results were observed under high correlation scenar-ios (*ρ* = 0.5, 0.7) and in presence of dependence between items, see Table 18-22 in Suppl. Mat.

In terms of computational performance, the average computational time for qch_copula with *Q* = 16 and *n* = 10^5^ was 78 minutes for the model-fitting step and approximately 1 minute per tested composite hypothesis. The analysis was conducted on a computing platform with one thread and 3.225 GB of allocated RAM, running under the Ubuntu Linux operating system.

### Application I: Detection of pleiotropic regions associated to psychiatric disorders

To illustrate our method, we performed a comprehensive analysis of 14 psychiatric disorders using data derived from genetic association studies conducted by the Psychiatric Genomics Consortium, see Section 7.A and Table 23 in Suppl. Mat. for details. The initial data consists of *p*-values obtained for *n* = 6, 267, 062 to 12, 438, 502 single nucleic polymorphisms (SNPs) for the different studies. The analysis aims to identify pleiotropic genomic regions simultaneously associated with multiple disorders. In the initial analysis of (20), the authors applied the MAGMA method (21) to aggregate *p*-values at the gene level before running their analysis, drastically reducing the size of the data to 26,024 genes. Aggregated data were analysed using PLACO (8) to detect genes associated with two or more disorders, focusing on the top genes detected in at least eight disorders. As the number of traits allowed by PLACO is limited to 2, all pairwise analyses of 2 among the 14 traits were performed. For a given gene, the list of associated disorders corresponded to those found significant in at least one pair-wise analysis.

The initial approach suffers from several technical limitations. Firstly, while the aggregation step reduces the computational burden of the CHT procedure, it may lead to significantly less accurate outcomes. This is because some regions (e.g. intergenic regions or genes not listed in the annotation list) may not be represented, but also because the detection resolution is now limited to the gene level. Second, the use of a pairwise approach may yield ambiguous or inconsistent results. For example, gene TMX2 was detected in the ADHD-AN and the AN-SCZ analyses but not in the ADHD-SCZ analysis. Similarly, gene MIR2113, one of the two genes “that are identified in 10 psychiatric disorders”, is declared significant in only 26 out of the 45 PLACO pairwise comparisons involving the ten candidate disorders, with Anorexia Nervosa (AN) being identified in only 1 out of its 13 associated pairwise comparisons. Finally, note that aggregating the results of all pairwise analyses does not come with any statistical guarantee regarding false positive control for the final gene list.

The goal of detecting pleiotropic SNPs exhibiting an effect in at least *q* disorders can be turned into a composite hypothesis testing procedure where the corresponding composite null hypothesis for SNP *i* is defined as follows:

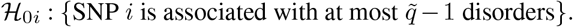

Following the initial analysis, we focused on 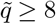 and performed CHT at a nominal type-I error rate *α* = 0.05. We ran qch_copula on the 5,172,884 SNPs common to all analyses and compared the outcomes with those obtained using the PLACO pairwise analyses. The results are displayed in Table 5 (see Suppl. Mat. Fig. 16 for quality control of the *p*-values distribution and Fig. 17 for the Manhattan plots).

**Table 5.** Number of candidate genes and SNPs identified by PLACO and qch_copula, respectively, for different levels of pleiotropy (i.e., associated with at least the specified number of disorders).

| At least # of disorders | 8 | 9 | 10 | 11 | >11 |
| --- | --- | --- | --- | --- | --- |
| PLACO | 38 | 21 | 2 | 0 | 0 |
| qch_copula | 1,608 | 498 | 211 | 25 | 0 |

Focusing on the composite hypothesis test with 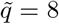, our method identified 1,608 SNPs (see Fig 2 for their description), corresponding to 28 distinct regions. The regions identified by qch_copula may include more than one gene. Out of the 38 candidate genes detected using the PLACO approach, 35 were also found by our method (i.e. at least one significant SNP within the gene was identified). The three remaining genes NEGR1, TMX2, and C11orf31 that were identified exclusively by PLACO had only 7, 4, and 3 *p*-values below 0.01 out of 14 obtained from the MAGMA analysis (for each gene), respectively. Since the goal was to detect genes associated with a minimum of eight disorders, one would expect the number of significant MAGMA *p*-values to be close to eight or higher.

**Fig. 2.**
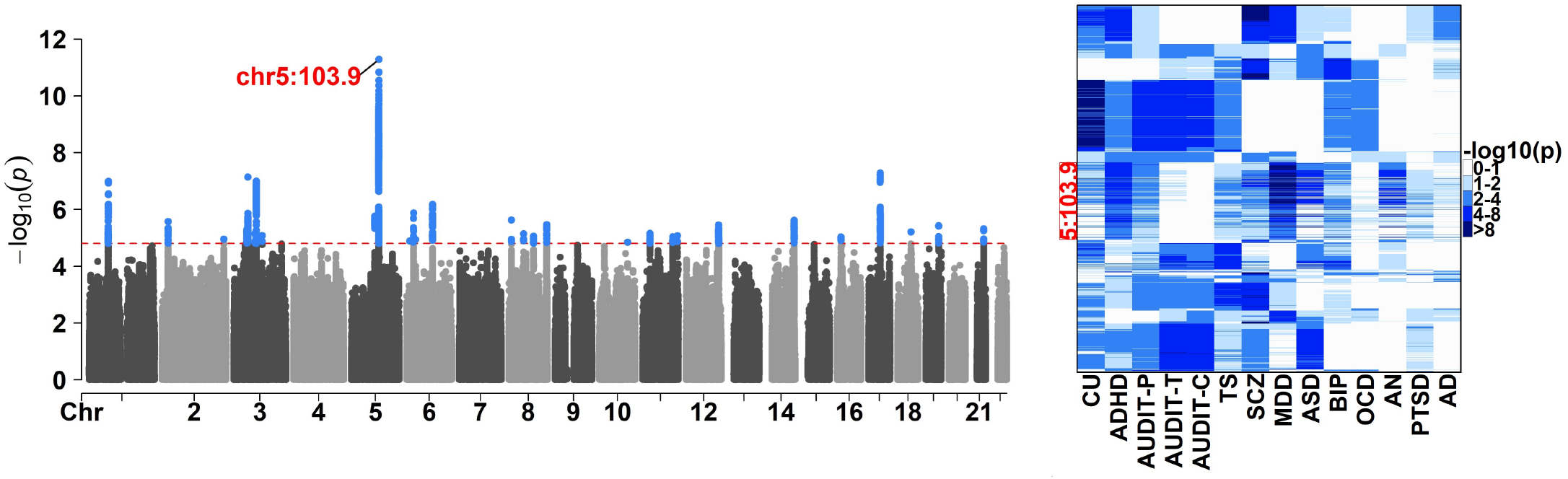
Results of the composite hypothesis test identifying SNPs associated with at least eight disorders. **Left:** -log10(*p*-values) of the composite hypothesis test along the chromosomes. The red dotted line represents the significance threshold at a nominal false discovery rate of 0.05. Significant SNPs are represented in blue. **Right:** Initial GWAS -log10(*p*-values) of the significant SNPs (in rows).

Additionally, qch_copula identified eight new regions, each including more than 10 SNPs, that were not detected by PLACO (see Table 6 for their description). Three of these eight regions were not detected in the initial analysis because the annotation list contained no gene located in these regions. Among them, the top region identified by qch_copula, located on chromosome 5 contained 338 SNPs (see the corresponding peak on the Manhattan plot in Fig 2, left). Notably, 25 of these SNPs were detected as associated with 11 disorders (see the corresponding initial GWAS -log10(*p*-values) in Fig 2, right). This region shares an overlap with the gene RP11-6N13.1, which has been recurrently reported to be associated with psychiatric disorders (22–26), including ADHD, ASD, BIP, MDD, SCZ and TS, among others.

**Table 6.** Summary information of the eight new regions associated with at least eight disorders identified by qch_copula.

| Chr | Position (Mb) | Nb SNPs | Top SNP | Pval top SNP |
| --- | --- | --- | --- | --- |
| 1 | 73.8 - 73.9 | 156 | rs11210259 | 1.013e-07 |
| **2** | **22.5 - 22.6** | 14 | rs11688810 | 2.665e-06 |
| 3 | 52.6 - 53.1 | 90 | rs11922961 | 7.273e-08 |
| **5** | **103.6 - 104.0** | 338 | rs30266 | 5.162e-12 |
| 6 | 25.9 - 26.3 | 12 | rs115810 | 4.773e-06 |
| 8 | 143.3 - 143.3 | 11 | rs4458978 | 3.416e-06 |
| **11** | **27.6 - 27.6** | 13 | rs11030084 | 6.959e-06 |
| 11 | 113.2 - 113.3 | 16 | rs2734837 | 9.245e-06 |

Note: The regions not included in the initial analysis due to the absence of any gene in the annotation list are denoted in boldface.

### Application II: Detection of viruses resistance hotspots regions in cucumber

Our second application case is based on GWAS summary statistics derived from the experiment described in (27). In this study, a panel of 226 cucumber elite lines, 40 landraces and 23 hybrids were inoculated with six viruses (denoted CGMMV, CMV, CVYV, PRSV, WMV and ZYMV hereafter) to evaluate their responses, see Section 7.B in Suppl. Mat. for details. GWAS were conducted on each of the 6 virus separately (refered to as individual GWAS hereafter), on a number of SNPs ranging from *n* = 378, 049 to *n* = 424, 393. The aim of the study was to identify QTLs associated with virus resistance in cucumber.

In the original analysis QTLs associated with resistance against multiple viruses were detected through a simple intersection of the lists of putative QTLs obtained from separate GWAS runs for each virus. As an alternative approach we addressed the detection of SNPs associated with resistance to at least 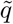 viruses through composite hypothesis testing. The corresponding null hypothesis for SNP *i* was defined as follows:

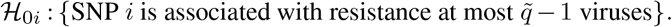

We focused on values 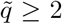 and applied the qch_copula method to the 339,804 SNPs common to all analyses, after excluding the genomic region spanning from 6.7 Mb to 12.9 Mb on chromosome 2, which corresponds to a non-recombinant region that produced anormalous *p*-value distributions. The nominal type-I error rate was set at *α* = 0.05. Our analysis identified 1,845, 164, and 15 SNPs associated with resistance to at least two, three, and four viruses, respectively (see Supplementary Materials, Fig. 19 for quality control of *p*-value distributions, and Fig. 20 for Manhattan plots). For 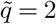, the 1,845 SNPs mapped to five distinct genomic regions on chromosomes 1, 2, 5, and 6 (see Table 7 for details). All significant SNPs had at least two *p*-values below 10^−4^ out of the six GWAS (Fig 3). Analysis of the initial GWAS *p*-values indicates that the significant regions on chromosomes 1, 2, and 6 are associated with resistance to PRSV and ZYMV. In contrast, the region on chromosome 5 is associated with resistance to the remaining four viruses: WMV, CGMMV, CVYV and CMV. Notably, this region was also significant when testing for association with at least three or four viruses - no SNP being associated with resistance to more than four viruses.

**Table 7.**
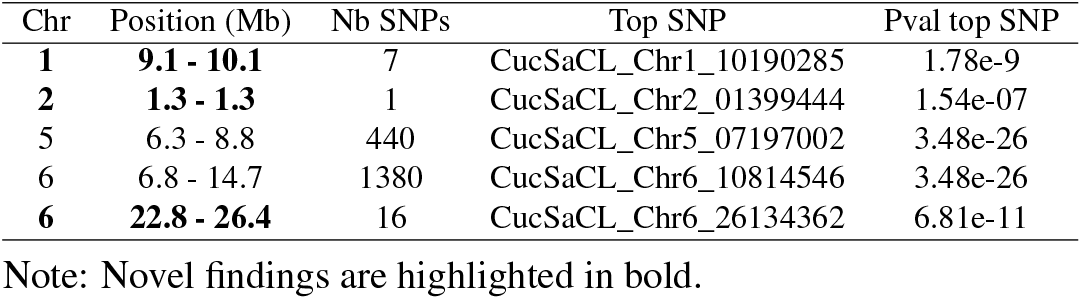
Summary of the five genomic regions associated with resistance to at least two viruses identified by qch_copula.

**Fig. 3.**
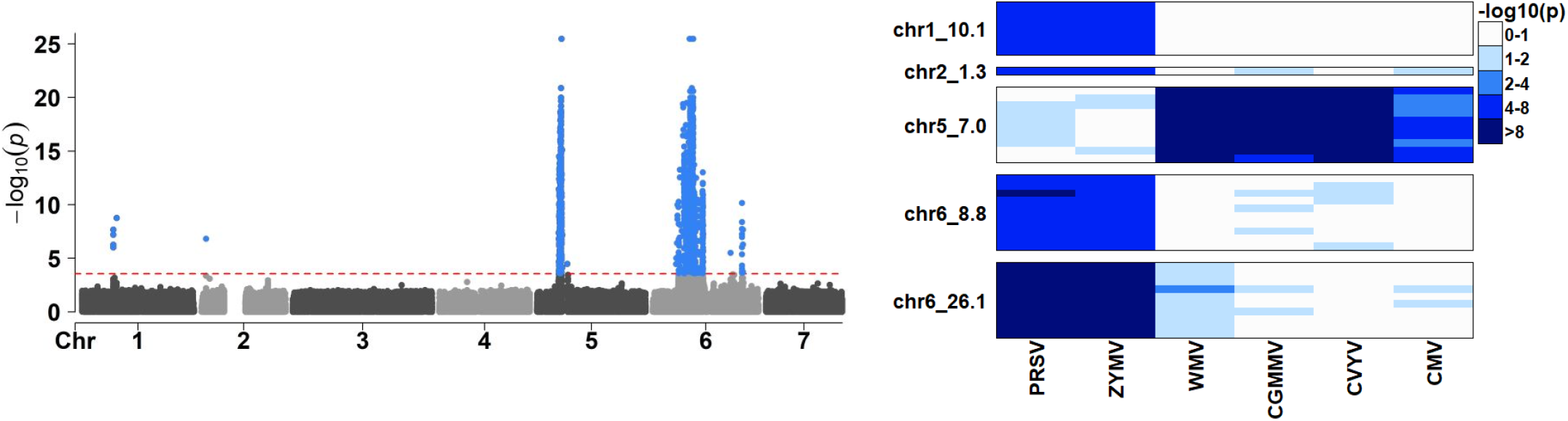
Results of the composite hypothesis test identifying SNPs associated with resistance to at least two viruses. **Left:** -log10(*p*-values) of the composite hypothesis test along the chromosomes. The red dotted line represents the significance threshold at a nominal false discovery rate of 0.05. Significant SNPs are represented in blue. **Right:** Initial GWAS -log10(*p*-values) of the top ten significant SNPs per regions identified (in rows).

Importantly, three of the detected hotspot regions were not reported in the original study (Table 7), of which two are strongly supported by external studies, confirming the hypothesis of a shared resistance mechanism across several viruses in these two regions. The hotspot region on chromosome 2 is associated with resistance to PRSV and ZYMV but also colocalizes with putative quantitative trait loci (QTLs) previously reported to confer resistance to CMV and CABYV (28). Similarly, the region on chromosome 6 (22.8–26.4 Mb) linked to resistance to PRSV and ZYMV, and colocalizes with QTLs associated with WMV (29) and CABYV (28). This highlights the enhanced statistical power of the joint analysis with a dedicated CHT procedure compared to individual GWAS approaches.

## Conclusion

We have developed a novel approach called qch_copula for composite hypothesis testing based on a multivariate mixture model. Our method comes with a rigorously defined *p*-value directly obtained from the mixture model approach, which, to our knowledge, is the first of its kind.

A key feature of qch_copula is its ability to flexibly model the conditional distributions, representing the probabilistic patterns by which each component of the mixture produces observable data. Previous published methods rely on fully parametric models, assuming both the 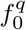 and 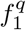 distributions of Equation Eq. (3) to be Gaussian density functions (7, 10), or (a mixture of) *χ*^2^ density functions, after data transformation (9). Although such modelings allow one to explicitly account for correlations between traits, they significantly constrain the shape of 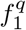. In comparison, qch_copula performs a nonparametric estimation of the alternative marginal distribution while still accounting for dependencies through the use of the copula function. This adaptive estimation procedure allows for a better fit of the data, resulting in efficient control of Type I errors and improved detection performance. In practice, the correlation matrix is inferred jointly with the prior proportions within the (M) step of the EM algorithm, enhancing our ability to capture the dependence more accurately compared to methods like Primo and PLACO, where the correlation matrix is estimated upstream.

Our model assumes that the copula function is the same across all components, employing a single correlation matrix to model the dependence structure. While this may be seen as a limitation, it leads to practical benefits. Modeling the dependency component-wise, as in the IMIX procedure, can lead to a prohibitive computational burden and downgrade the results. This is because some components may be poorly represented in the dataset, compromising the inference of the associated correlation matrices. Alternatively, ignoring dependencies leads to significantly inflated Type I error rates, as highlighted by the performance of qch (without copula) in our simulation experiment. The qch_copula procedure strikes a favorable balance between data fitting and computational efficiency in the inference procedure, offering a more practical and effective solution while capturing the essential structure of dependence among the multivariate p-values.

Another hypothesis shared by most, if not all, CHT procedures is the independence between the z-scores within each series, a condition that may be unrealistic in many applications, such as genomics, where test statistics often exhibit local dependence. Our simulation study demonstrated that a moderate within-series correlation (*ξ* = 0.3) has minimal impact on the overall performance of most procedures, and the general conclusions regarding the performance of the procedures remained unchanged. Interestingly, when dependence between items was introduced, the qch_copula method showed improved FDR control, particularly in scenarios where slight inflation had been observed under the independence assumption (e.g., *Q* = 16, *ρ* = 0.3, testing at least 16 traits; see Tables 3 and 4). These results suggest that qch_copula is not only designed to accommodate dependence across conditions but also demonstrates robustness to moderate item-level dependence. Dependencies between test statistics can be further addressed by applying multiple testing procedures that account for dependencies (30), or employing local score techniques that leverage, e.g., the spatial distribution of p-values throughout the genome to enhance detection power (31). Such methodologies are readily applicable to procedures that provide p-values, which is a key feature of our procedure.

Our simulation study highlights the challenges in achieving a balance between Type I error control (T1E) and detection power in the presence of correlation across varying values of *Q* (2, 8, 16). Notably, most methods struggled to maintain adequate FDR control across all scenarios while still achieving meaningful power, even when they explicitly model dependencies between test statistics. In the *Q* = 2 case, qch_copula was the only method that both controlled FDR and maintained a nonzero detection power although moderate.

For larger values of *Q*, qch_copula, Primo and adaFilter methods controlled FDR at the requested nominal level when the correlation between traits was low to moderate (i.e. *ρ* = 0 and *ρ* = 0.3), but the last two methods exhibited low power levels for some testing hypotheses. At higher correlation levels (*ρ* = 0.5 and *ρ* = 0.7), FDR inflation was observed for qch_copula, but these deviations remained more contained than those observed for Primo and adaFilter. It is worth mentioning that correlations higher than or equal to 0.5 can already be considered extreme values: in our two application cases, the empirical off-diagonal elements of the estimated copula correlation matrix were mostly below 0.09 and 0.12, respectively (90th percentiles), as shown in Fig. 15 and 18 in Supp. Mat.

The two CHT applications on real data requested the use of a null composite hypothesis of the form “item *i* has an effect in at most 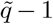 studies/traits”. The choice of 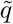 was suggested by previous analyzes (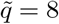 in Application I) or by the hypothesis to be tested (pleiotropy, i.e. 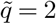 in Application II) but alternative values of 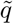 were also considered. Considering several values for 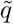 may be a simple way to rank the items - in the present case, the higher the value of 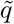 for which the composite hypothesis is rejected the better. Alternatively, the value of 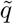 may be set based on considerations about budget and expected gains: in the resistance to pathogen case, the choice of 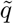 can be based on on the (financial) added value a resistance to 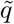 pathogens would confer to new cultivars compared to existing ones. Considering more than one value for 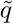 raises the question of post hoc inference, i.e. of the a posteriori choice of 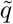 once the data are observed (32), which is a consideration for future work.

Our procedure relies on summary statistics (i.e. the z-scores computed from the *p*-values of the initial test series). As such, it belongs to the larger family of meta-analysis procedures that have been developed in the last decade in the context of genetic association.

Most existing multi-trait methods based on summary statistics (33–39) focus on testing the overall mean or variance of allelic effects over the traits, which corresponds to the special case of testing the association with at least one trait in our CHT setting. However,

CHT allows testing a much wider range of composite hypotheses, as any two complementary sets of configurations can be defined for ℋ_0_ and ℋ_1_. It is important to note that different series of composite hypotheses may be tested in parallel since the testing step stands independently of the model inference step and does not require parameter re-estimation. Importantly, relying on series of *p*-values exclusively makes our procedure amenable to data integration of studies with diverse outcomes, including continuous, binary, and time-to-event measurements. The exclusive reliance on *p*-values is also particularly advantageous when direct access to the raw data is constrained due to ethical considerations or data confidentiality.

In certain applications, it is crucial to account for the direction of effects for e.g. identifying markers that influence multiple traits or diseases in a consistent direction. This requires incorporating effect signs into the statistical model. An extension of our composite hypothesis testing procedure accounting for the direction of effects in the conditionally independent case is readily available in the qch package. Extending the performance to account for both sign effects and dependencies between series will be the subject of future work.

The proposed methodology is implemented in the R package `qch`, which is publicly available on CRAN.

## ACKNOWLEDGEMENTS

The authors would like to thank the Psychiatric Genomics Consortium (PGC) for making the summary statistics of psychiatric disorders GWAS publicly available. Additionally, the authors sincerely thank Séverine Monnot for sharing the summary statistics of the cucumber virus GWAS and for her valuable assistance in interpreting the results of our analysis. The authors are grateful to the INRAE MIGALE bioinformatics facility (MIGALE, INRAE, 2020. Migale bioinformatics Facility, doi: 10.15454/1.5572390655343293E12) for providing computing resources.

## DATA AVAILABILITY

The psychiatric disorders GWAS summary statistics are available in the Psychiatric Genomics Consortium repository at https://pgc.unc.edu/for-researchers/download-results/, and can be accessed with the identifiers: an2017, anx2016, adhd2019, asd2019, sud2019-alcuse, bip2019, mdd2018, ocd2018, ptsd2019, scz2019asi and ts2019. The cucumber viruses resistance GWAS summary statistics are available in the following data repository: https://doi.org/10.57745/XQ3P72. Functions to perform composite hypothesis procedure can be found in the R package qch, available at https://cran.r-project.org/web/packages/qch/index.html. All the codes used for the different analyses presented in the article are available in the following public Git repository : https://forge.inrae.fr/annaig.de-walsche/qch_copula_article_code.git.

## Supplementary Note 1: Estimation of the proportions *w*_*c*_ and the copula parameter *θ* using an EM algorithm

### Initialisation

The prior proportions are initialized using

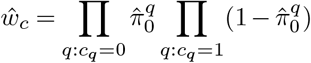

where 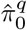 corresponds to the marginal prior probability in model (3) estimated in the first step of the inference procedure. Parameter *θ* is initialized using the inversion of Kendall’s tau estimator (40):

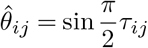

where *τ*_*ij*_ is the Kendall’s tau for the random vector (*Z*_*i*_, *Z*_*j*_)^*T*^.

### Maximum likelihood estimate for the Gaussian copula parameter

The complete likelihood of the observations 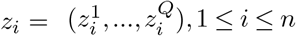 and the latent variables 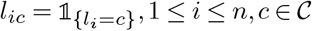 is given by:

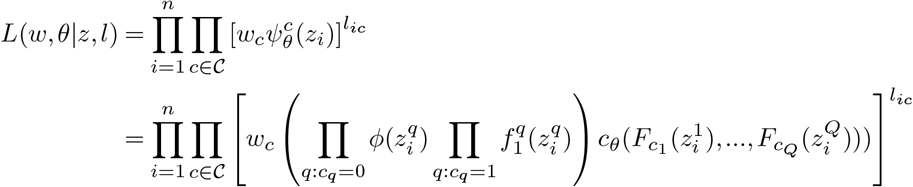

Define *Q* (*w, θ*|*w*^(*t*)^, *θ*^(*t*)^) as the expected value of the complete log-likelihood, with respect to the current conditional distribution of *L* given *Z* and the current estimates of the parameters *w*^(*t*)^ and *θ*^(*t*)^:

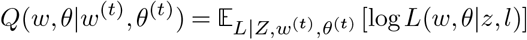

We denote 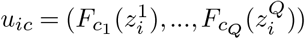 and 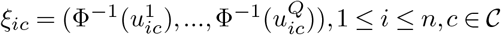. One has:

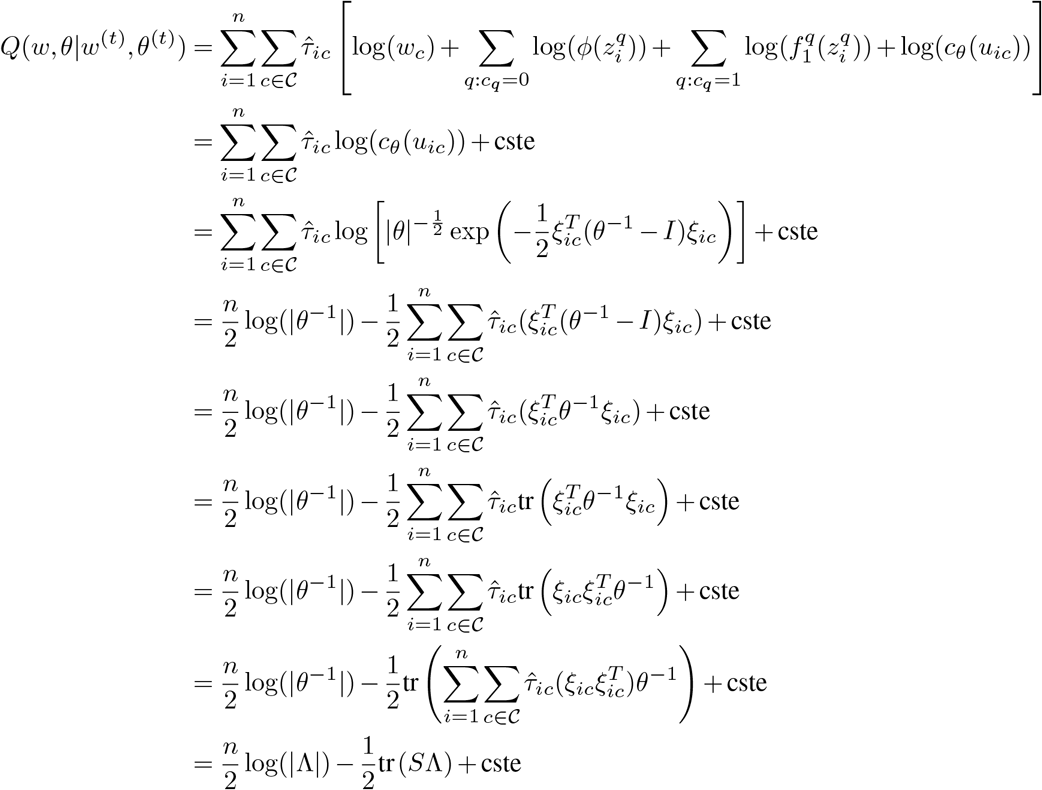

where 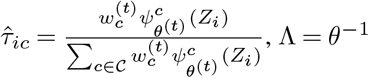 and 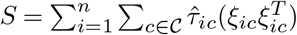.

Differentiating the log-likelihood with respect to Λ = *θ*^−1^ leads to

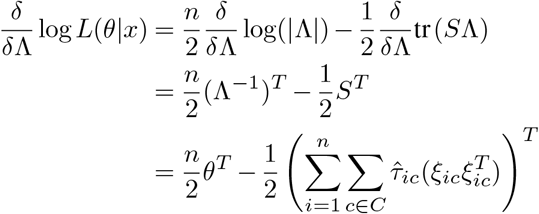

Setting the derivative at 0, one obtains

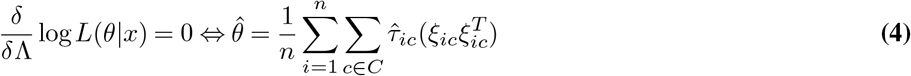

Note that the explicit expression of 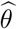 obtained in Eq. (4) guarantees that the estimated matrix is symmetric positive-definite, but not necessarily with value 1 on the diagonal. In practice, one needs to check that 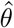 is properly a correlation matrix at each (M) step of the EM procedure, and whenever needed correct the initial estimate by scaling the resulting covariance matrix into a correlation matrix.

## Supplementary Note 2: Equivalence between Benjamini-Hochberg correction on p-values and the estimation of lFDR from the posteriors

In the classical setting of the test procedure, one can control the FDR at a nominal level of *α* by applying the Benjamini-Hochberg (BH) procedure. Assume *n* test were performed and let *p*_(1)_ ≤ … ≤*p*_(*n*)_ be the set of ordered p-values. Define the value *r* such as

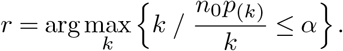

The set of rejected hypotheses is then *S*_*BH*_ = {*i*_(1)_, …, *i*_(*r*)_ }, where *i*_(*k*)_ is the index associated to p-value *p*_(*k*)_. Here *n*_0_ is the number of items for which ℋ_0_ holds, which is usually unknown. In practice, *n*_0_ can be upper bounded by *n* or adaptively obtained from the data, which will be designated hereafter by the adaptive BH procedure(16).

Alternatively, in the setting of a mixture model with two components 0 and 1 with emission distributions *ψ*_0_, *ψ*_1_ and priors *W*_0_, *W*_1_, one can define the posterior

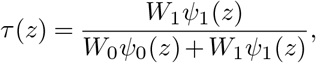

The local FDR can then be approximately controlled at a nominal level of *α* by applying the following empirical procedure (12, 13). Assume *n* items were generated under the previous mixture model and let *τ*_(1)_ ≤… ≤*τ*_(*n*)_ be the set of ordered posteriors. Define the value *m* such as

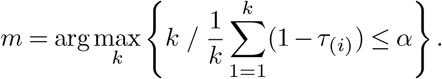

The set of items classified into class 1 is then *S*_*lF DR*_ = {*i*_(1)_, …, *i*_(*m*)_}, where *i*_(*k*)_ is the index associated to posterior *τ*_(*k*)_. The connection between the two approaches in the context of CHT is established by the next proposition:

### Proposition 1

Consider *n* observations drawn according to mixture model Equation 1 of Section Model, and let 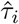 and 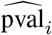 be the posterior and p-value associated to item *i*, respectively. Consider the adaptive BH procedure where *n*_0_ is estimated by *nŴ*_0_, where 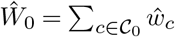. Then *S*_*BH*_ = *S*_*lF DR*_.

*Proof:*

For a given item *i*, the p-value corresponding to the composite hypothesis test is constructed as follows:

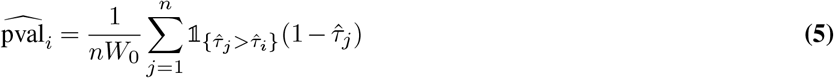

Let 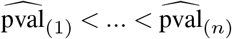 represent the ordered set of p-values, and *i*_(*k*)_ is the index associated to p-value 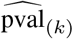.

According to the adaptive BH procedure, the estimate of the False Discovery Rate when the set of rejected hypotheses is *S*_*BH*_ = {*i*_(1)_, …, *i*_(*k*)_} is given by:

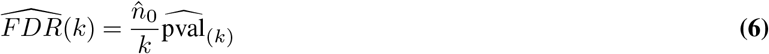

with 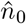 an estimate of the number of true null hypotheses. Here, one can estimate the number of null hypotheses as 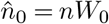. Hence, we have:

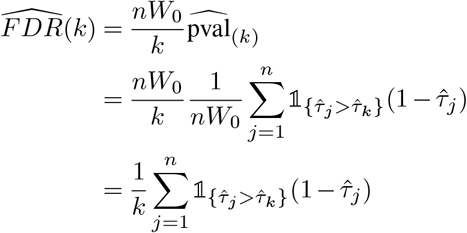

By the definition of the p-value 5, 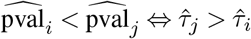.

Consequently:

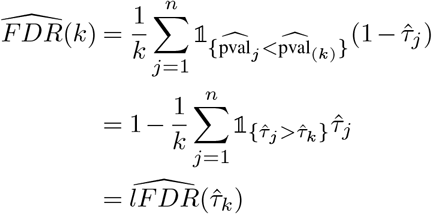

where 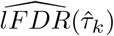 is the estimate of the false discovery rate when thresholding at the level 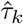 defined by (12) and (13). Thus, applying the adaptive BH correction with 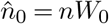 to p-value Eq. (5) is equivalent to using the local FDR rejection rule defined by (12) and (13). ▪

## Supplementary Note 3: Algorithm for the Memory-Efficient EM

Here we present how the burden associated to the storage of the posterior matrix can be avoided in the EM algorithm. Quantities in red correspond to the ones that are actually stored in place of the posterior matrix for later use.

### Algorithm 1 Proportions *w*_*c*_ and copula parameter *θ* inference using EM with memory handling

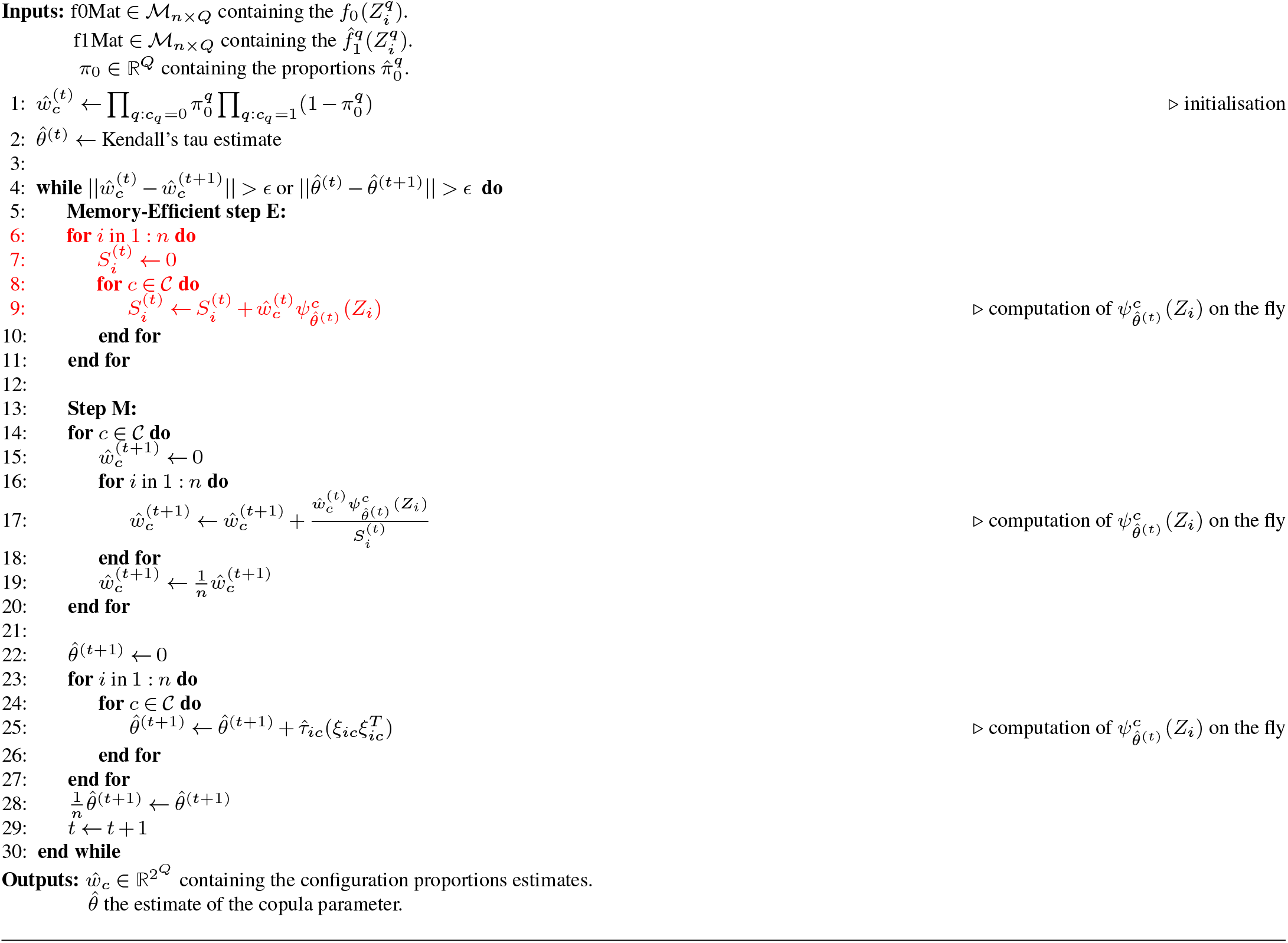

## Supplementary Note 4: Details for generating each simulation setting

**Table 8.** Description of the 48 settings considered for the simulation study.

| Spatial correlation ($\xi$) | $Q$ | Correlation ($\rho$) | Scenario | Setting |
| --- | --- | --- | --- | --- |
| 0 | 2 | 0 | sparse | 1 |
|  |  |  | dense | 2 |
|  |  | 0.3 | sparse | 3 |
|  |  |  | dense | 4 |
|  |  | 0.5 | sparse | 5 |
|  |  |  | dense | 6 |
|  |  | 0.7 | sparse | 7 |
|  |  |  | dense | 8 |
|  | 8 | 0 | sparse | 9 |
|  |  |  | dense | 10 |
|  |  | 0.3 | sparse | 11 |
|  |  |  | dense | 12 |
|  |  | 0.5 | sparse | 13 |
|  |  |  | dense | 14 |
|  |  | 0.7 | sparse | 15 |
|  |  |  | dense | 16 |
|  | 16 | 0 | sparse | 17 |
|  |  |  | dense | 18 |
|  |  | 0.3 | sparse | 19 |
|  |  |  | dense | 20 |
|  |  | 0.5 | sparse | 21 |
|  |  |  | dense | 22 |
|  |  | 0.7 | sparse | 23 |
|  |  |  | dense | 24 |
| 0.3 | 2 | 0 | sparse | 1s |
|  |  |  | dense | 2s |
|  |  | 0.3 | sparse | 3s |
|  |  |  | dense | 4s |
|  |  | 0.5 | sparse | 5s |
|  |  |  | dense | 6s |
|  |  | 0.7 | sparse | 7s |
|  |  |  | dense | 8s |
|  | 8 | 0 | sparse | 9s |
|  |  |  | dense | 10s |
|  |  | 0.3 | sparse | 11s |
|  |  |  | dense | 12s |
|  |  | 0.5 | sparse | 13s |
|  |  |  | dense | 14s |
|  |  | 0.7 | sparse | 15s |
|  |  |  | dense | 16s |
|  | 16 | 0 | sparse | 17s |
|  |  |  | dense | 18s |
|  |  | 0.3 | sparse | 19s |
|  |  |  | dense | 20s |
|  |  | 0.5 | sparse | 21s |
|  |  |  | dense | 22s |
|  |  | 0.7 | sparse | 23s |
|  |  |  | dense | 24s |

### Independence between markers within series

Each dataset consists of a *n × Q* matrix of Z-scores generated as follows. Markers are clustered into *K* + 1 clusters. Within each cluster *k*, all markers share the same configuration vector *c*_*k*_ = (*c*_*k*1_, …, *c*_*kQ*_) ∈ {0, 1 }^*Q*^, indicating the presence (1) or absence (0) of association for each traits. The z-scores of markers in cluster *k* are independently generated according to:

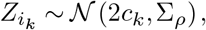

where Σ_*ρ*_ is a *Q × Q* correlation matrix where all covariances between traits are set to *ρ*. The covariance structure Σ_*ρ*_ is common across all clusters. Hence, a cluster is fully specified by its configuration vector and the number of markers it contains.

### Dependence between markers within series

To account for dependence between markers within each series, we extend the previous model by introducing correlation between the markers within each cluster. As before, markers are organized into *K* + 1 clusters, where each cluster *k* is defined by a configuration vector *c*_*k*_ = (*c*_*k*1_, …, *c*_*kQ*_) and a number of markers *n*_*k*_. For each cluster *k*, we generate an *n*_*k*_ *× Q* matrix *Z*_*k*_, corresponding to the z-scores of the *n*_*k*_ markers across the *Q* traits. This matrix is drawn according to:

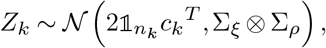

where Σ_*ξ*_ is a *n*_*k*_ *× n*_*k*_ correlation matrix with all off-diagonal entries equal to *ξ*, modelling the correlation between markers and Σ_*ρ*_ is a *Q×Q* correlation matrix with all off-diagonal entries equal to *ρ*, modeling correlation between traits.

To mimic a more realistic genomic structure, large clusters of markers corresponding to the global null configuration (i.e., *c* = (0, …, 0)) are further subdivided into smaller clusters of size 50, thereby introducing local dependence.

We considered different values for *Q* (2,8,16), two scenarios: *sparse* or *dense*, and four levels of correlation (*ρ* = 0, 0.3, 0.5, 0.7). When applicable, the spatial correlation *ξ* was set to 0.3.

### Details of clusters for all settings

Denoting 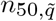 and 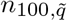 the number of clusters of 50 and 100 markers respectively, which are associated with exactly 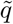 randomly chosen traits, the details for generating a dataset for each simulation setting are presented below. Additionally, a final cluster is included to account for all remaining markers; this cluster follows the complete null configuration, i.e., *c* = (0, 0,…, 0).

#### *Settings 1 to 8: Q* = 2

- Sparse scenario: *n*_50,1_ = *n*_50,2_ = 20 and *n*_100,1_ = *n*_100,2_ = 10, such that *w*_00_ = 0.96, *w*_01_ + *w*_10_ = 0.02 and *w*_11_ = 0.02.
- Dense scenario: *n*_50,1_ = 400, *n*_100,1_ = 200 and *n*_50,2_ = 100, *n*_100,2_ = 50, such that *w*_00_ = 0.5, *w*_01_ + *w*_10_ = 0.4 and *w*_11_ = 0.1.

#### *Settings 9 to 16: Q* = 8

- Sparse scenario: *n*_50,*j*_ = 20 and *n*_100,*j*_ = 10 for *j* ∈ {1, 2, 4, 6, 8}, such that *w*_0_ = 0.90 for the markers being associated with none of the traits and *w*_*j*_ = 0.02 for the markers being associated with exactly *j* ∈ {1, 2, 4, 6, 8} traits.
- Dense scenario: *n*_50,*j*_ = 100 and *n*_100,*j*_ = 50 for *j* ∈ {1, 2, 4, 6, 8}, such that *w*_0_ = 0.50 for the markers being associated with none of the traits and *w*_*j*_ = 0.1 for the markers being associated with exactly *j* ∈ {1, 2, 4, 6, 8} traits.

#### *Settings 17 to 24: Q* = 16

- Sparse scenario: *n*_50,*q*_ = 20 and *n*_100,*q*_ = 10 for 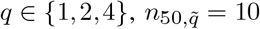 and 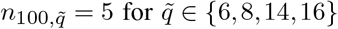, such that *w*_0_ = 0.90 for the markers being associated with none of the traits, *w*_*q*_ = 0.02 for the markers being associated with exactly *q* ∈ {1, 2, 4} traits, and 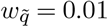 for the markers being associated with exactly 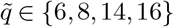 traits.
- Dense scenario: *n*_50,*q*_ = 100 and *n*_100,*q*_ = 50 for 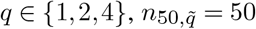 and 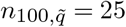 for 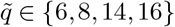, such that *w*_0_ = 0.5 for the markers being associated with none of the traits, *w*_*q*_ = 0.1 for the markers being associated with exactly *q* ∈ {1, 2, 4} traits, and 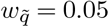 for the markers being associated with exactly 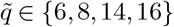 traits.

## Supplementary Note 5: Additional simulations results

**Table 9.**
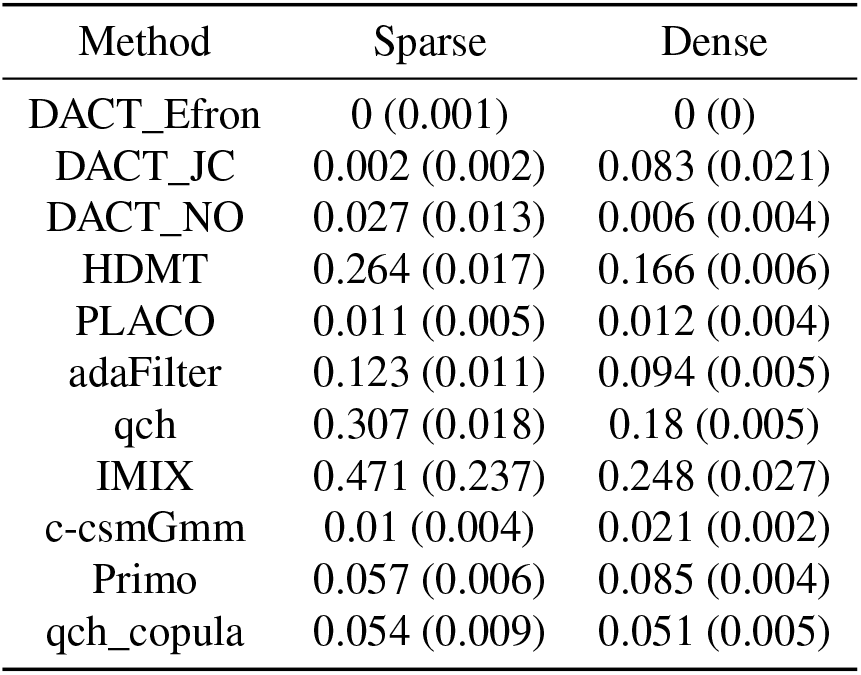
Detection power of the procedures for sparse and dense scenarios with *Q* = 2, *ρ* = 0.3 and *ξ* = 0 (settings 1 and 2).

| Method | Sparse | Dense |
| --- | --- | --- |
| DACT_Efron | 0 (0.001) | 0 (0) |
| DACT_JC | 0.002 (0.002) | 0.083 (0.021) |
| DACT_NO | 0.027 (0.013) | 0.006 (0.004) |
| HDMT | 0.264 (0.017) | 0.166 (0.006) |
| PLACO | 0.011 (0.005) | 0.012 (0.004) |
| adaFilter | 0.123 (0.011) | 0.094 (0.005) |
| qch | 0.307 (0.018) | 0.18 (0.005) |
| IMIX | 0.471 (0.237) | 0.248 (0.027) |
| c-csmGmm | 0.01 (0.004) | 0.021 (0.002) |
| Primo | 0.057 (0.006) | 0.085 (0.004) |
| qch_copula | 0.054 (0.009) | 0.051 (0.005) |

For each procedure the detection power (averaged over 25 runs) are displayed. Numbers in brackets correspond to standard errors.

**Table 10.** False discovery rate and detection power of the procedures in sparse and dense scenarios with *Q* = 2, *ρ* = 0 and *ξ* = 0 (settings 1 and 2).

| Method | Sparse |  | Dense |  |
| --- | --- | --- | --- | --- |
|  | FDR | Power | FDR | Power |
| DACT_Efron | 0.002 (0.013) | 0.001 (0.002) | 0.004 (0.01) | 0.003 (0.005) |
| DACT_JC | 0.023 (0.012) | 0.064 (0.021) | **0.248 (0.014)** | 0.408 (0.024) |
| DACT_NO | 0.002 (0.013) | 0.001 (0.002) | 0 (0) | 0 (0) |
| HDMT | 0.026 (0.014) | 0.071 (0.019) | 0.053 (0.011) | 0.051 (0.007) |
| PLACO | 0.047 (0.018) | 0.065 (0.007) | **0.079 (0.02)** | 0.022 (0.005) |
| adaFilter | 0.02 (0.013) | 0.051 (0.012) | 0.039 (0.011) | 0.026 (0.005) |
| qch | 0.035 (0.01) | 0.112 (0.01) | 0.05 (0.011) | 0.052 (0.004) |
| IMIX | 0.056 (0.017) | 0.152 (0.013) | 0.048 (0.011) | 0.055 (0.012) |
| c-csmGmm | **0.076 (0.042)** | 0.034 (0.009) | 0.055 (0.015) | 0.015 (0.002) |
| Primo | 0.016 (0.012) | 0.049 (0.006) | 0.035 (0.013) | 0.022 (0.003) |
| qch_copula | 0.034 (0.011) | 0.107 (0.01) | 0.045 (0.01) | 0.043 (0.004) |

For each procedure the FDR and the detection power (averaged over 25 runs) are displayed. Numbers in brackets correspond to standard errors. False discovery rate above 0.05 are in boldface.

**Table 11.** False discovery rate and detection power of the procedures in sparse and dense scenarios with *Q* = 2, *ρ* = 0.5 and *ξ* = 0 (settings 5 and 6).

| Method | Sparse |  | Dense |  |
| --- | --- | --- | --- | --- |
|  | FDR | Power | FDR | Power |
| DACT_Efron | 0 (0) | 0 (0) | 0 (0) | 0 (0) |
| DACT_JC | 0 (0) | 0 (0.001) | 0.026 (0.018) | 0.01 (0.006) |
| DACT_NO | **0.104 (0.033)** | 0.092 (0.022) | 0.055 (0.013) | 0.042 (0.008) |
| HDMT | **0.41 (0.017)** | 0.34 (0.011) | **0.374 (0.015)** | 0.53 (0.021) |
| PLACO | **0.3 (0.083)** | 0.007 (0.002) | 0.052 (0.025) | 0.011 (0.003) |
| adaFilter | **0.217 (0.02)** | 0.185 (0.014) | **0.121 (0.007)** | 0.146 (0.006) |
| qch | **0.437 (0.015)** | 0.387 (0.013) | **0.239 (0.006)** | 0.299 (0.006) |
| IMIX | **0.871 (0.021)** | 0.848 (0.063) | **0.295 (0.079)** | 0.448 (0.081) |
| c-csmGmm | **0.13 (0.176)** | 0.007 (0.003) | 0.057 (0.013) | 0.029 (0.003) |
| Primo | **0.081 (0.022)** | 0.057 (0.005) | **0.112 (0.009)** | 0.139 (0.005) |
| qch_copula | 0.057 (0.025) | 0.04 (0.007) | **0.061 (0.008)** | 0.067 (0.005) |

For each procedure the FDR and the detection power (averaged over 25 runs) are displayed. Numbers in brackets correspond to standard errors. False discovery rate above 0.05 are in boldface.

**Table 12.** False discovery rate and detection power of the procedures in sparse and dense scenarios with *Q* = 2, *ρ* = 0.7 and *ξ* = 0 (settings 7 and 8).

| Method | Sparse |  | Dense |  |
| --- | --- | --- | --- | --- |
|  | FDR | Power | FDR | Power |
| DACT_Efron | 0 (0) | 0 (0) | 0 (0) | 0 (0) |
| DACT_JC | 0 (0) | 0 (0) | 0.005 (0.021) | 0.001 (0.002) |
| DACT_NO | **0.255 (0.051)** | 0.182 (0.041) | **0.094 (0.011)** | 0.1 (0.009) |
| HDMT | **0.564 (0.011)** | 0.433 (0.013) | **0.459 (0.005)** | 0.637 (0.006) |
| PLACO | **0.663 (0.065)** | 0.01 (0.004) | **0.066 (0.027)** | 0.011 (0.003) |
| adaFilter | **0.379 (0.018)** | 0.273 (0.012) | **0.146 (0.008)** | 0.209 (0.006) |
| qch | **0.571 (0.017)** | 0.437 (0.018) | **0.352 (0.013)** | 0.428 (0.014) |
| IMIX | **0.915 (0.004)** | 0.87 (0.053) | **0.356 (0.015)** | 0.593 (0.021) |
| c-csmGmm | **0.072 (0.199)** | 0.006 (0.002) | **0.061 (0.009)** | 0.049 (0.004) |
| Primo | **0.113 (0.032)** | 0.051 (0.006) | **0.144 (0.008)** | 0.188 (0.005) |
| qch_copula | **0.063 (0.036)** | 0.03 (0.007) | 0.058 (0.008) | 0.107 (0.005) |

For each procedure the FDR and the detection power (averaged over 25 runs) are displayed. Numbers in brackets correspond to standard errors. False discovery rate above 0.05 are in boldface.

**Table 13.** False discovery rate and detection power of the procedures in sparse and dense scenarios with *Q* = 2, *ρ* = 0 and *ξ* = 0.3 (settings 1s and 2s).

| Method | Sparse |  | Dense |  |
| --- | --- | --- | --- | --- |
|  | FDR | Power | FDR | Power |
| DACT_Efron | 0.001 (0.003) | 0.003 (0.007) | 0.01 (0.015) | 0.006 (0.012) |
| DACT_JC | 0.021 (0.023) | 0.051 (0.048) | **0.25 (0.039)** | 0.403 (0.094) |
| DACT_NO | 0 (0.002) | 0.004 (0.011) | 0 (0) | 0 (0) |
| HDMT | 0.033 (0.024) | 0.081 (0.042) | 0.053 (0.015) | 0.053 (0.01) |
| PLACO | 0.056 (0.035) | 0.058 (0.033) | **0.066 (0.02)** | 0.02 (0.008) |
| adaFilter | 0.029 (0.029) | 0.041 (0.031) | 0.034 (0.014) | 0.025 (0.009) |
| qch | 0.033 (0.028) | 0.093 (0.058) | 0.054 (0.017) | 0.052 (0.011) |
| IMIX | 0.052 (0.012) | 0.178 (0.034) | 0.05 (0.01) | 0.058 (0.016) |
| c-csmGmm | **0.082 (0.047)** | 0.034 (0.02) | **0.068 (0.028)** | 0.014 (0.006) |
| Primo | 0.021 (0.029) | 0.043 (0.02) | 0.031 (0.015) | 0.022 (0.007) |
| qch_copula | 0.036 (0.031) | 0.084 (0.05) | 0.041 (0.012) | 0.036 (0.006) |

For each procedure the FDR and the detection power (averaged over 25 runs) are displayed. Numbers in brackets correspond to standard errors. False discovery rate above 0.05 are in boldface.

**Table 14.** False discovery rate and detection power of the procedures in sparse and dense scenarios with *Q* = 2 and *ρ* = 0.3 and *ξ* = 0.3 (settings 3s and 4s).

| Method | Sparse |  | Dense |  |
| --- | --- | --- | --- | --- |
|  | FDR | Power | FDR | Power |
| DACT_Efron | 0 (0) | 0 (0.001) | 0 (0) | 0 (0.001) |
| DACT_JC | 0.006 (0.02) | 0.007 (0.015) | **0.096 (0.03)** | 0.109 (0.053) |
| DACT_NO | 0.029 (0.038) | 0.033 (0.032) | 0.02 (0.016) | 0.01 (0.009) |
| HDMT | **0.242 (0.021)** | 0.27 (0.039) | **0.136 (0.016)** | 0.166 (0.024) |
| PLACO | **0.108 (0.099)** | 0.01 (0.008) | **0.068 (0.025)** | 0.016 (0.008) |
| adaFilter | **0.098 (0.027)** | 0.121 (0.032) | **0.086 (0.016)** | 0.094 (0.013) |
| qch | **0.261 (0.035)** | 0.311 (0.04) | **0.14 (0.016)** | 0.175 (0.019) |
| IMIX | **0.594 (0.156)** | 0.405 (0.291) | **0.181 (0.021)** | 0.238 (0.031) |
| c-csmGmm | **0.126 (0.105)** | 0.011 (0.008) | **0.062 (0.019)** | 0.021 (0.008) |
| Primo | 0.047 (0.028) | 0.058 (0.015) | **0.081 (0.016)** | 0.087 (0.01) |
| qch_copula | 0.038 (0.033) | 0.055 (0.031) | **0.062 (0.013)** | 0.056 (0.009) |

For each procedure the FDR and the detection power (averaged over 25 runs) are displayed. Numbers in brackets correspond to standard errors. False discovery rate above 0.05 are in boldface.

**Table 15.**
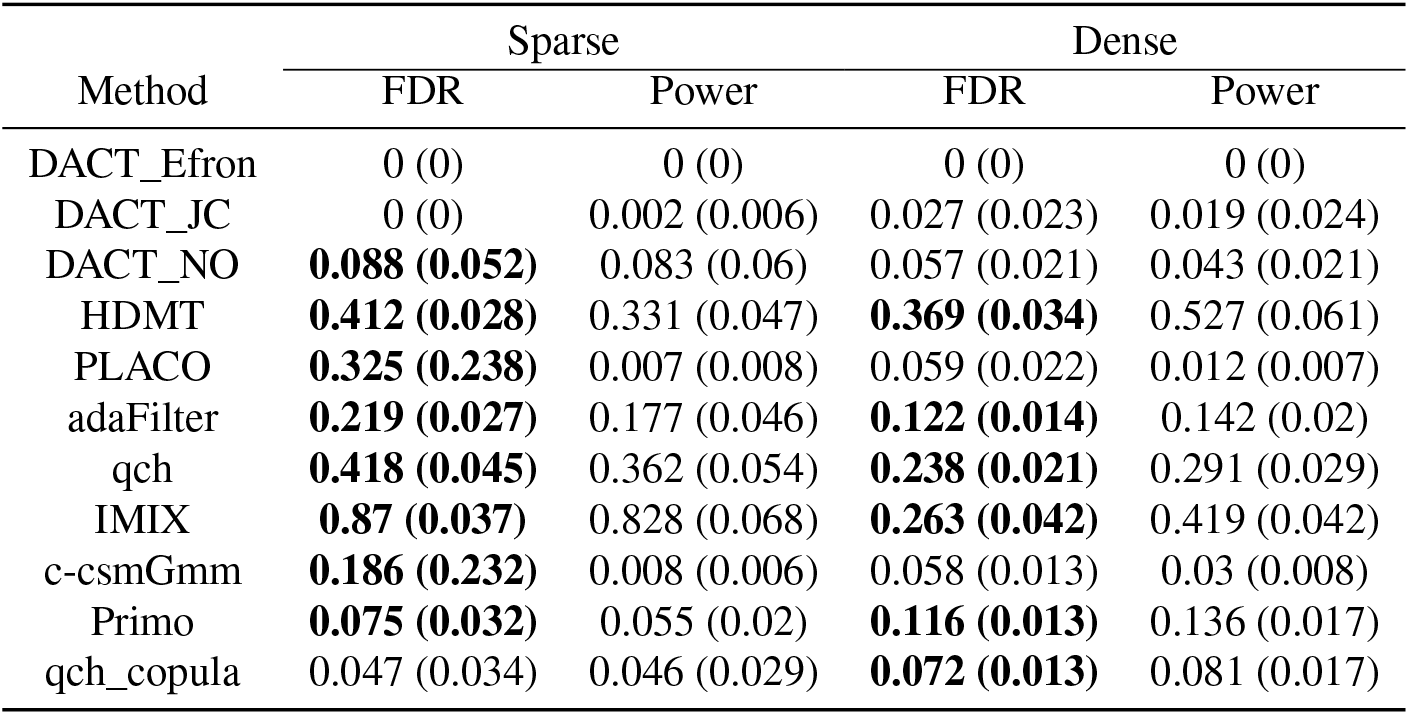
False discovery rate and detection power of the procedures in sparse and dense scenarios with *Q* = 2, *ρ* = 0.5 and *ξ* = 0.3 (settings 5s and 6s).

| Method | Sparse |  | Dense |  |
| --- | --- | --- | --- | --- |
|  | FDR | Power | FDR | Power |
| DACT_Efron | 0 (0) | 0 (0) | 0 (0) | 0 (0) |
| DACT_JC | 0 (0) | 0.002 (0.006) | 0.027 (0.023) | 0.019 (0.024) |
| DACT_NO | **0.088 (0.052)** | 0.083 (0.06) | 0.057 (0.021) | 0.043 (0.021) |
| HDMT | **0.412 (0.028)** | 0.331 (0.047) | **0.369 (0.034)** | 0.527 (0.061) |
| PLACO | **0.325 (0.238)** | 0.007 (0.008) | 0.059 (0.022) | 0.012 (0.007) |
| adaFilter | **0.219 (0.027)** | 0.177 (0.046) | **0.122 (0.014)** | 0.142 (0.02) |
| qch | **0.418 (0.045)** | 0.362 (0.054) | **0.238 (0.021)** | 0.291 (0.029) |
| IMIX | **0.87 (0.037)** | 0.828 (0.068) | **0.263 (0.042)** | 0.419 (0.042) |
| c-csmGmm | **0.186 (0.232)** | 0.008 (0.006) | 0.058 (0.013) | 0.03 (0.008) |
| Primo | **0.075 (0.032)** | 0.055 (0.02) | **0.116 (0.013)** | 0.136 (0.017) |
| qch_copula | 0.047 (0.034) | 0.046 (0.029) | **0.072 (0.013)** | 0.081 (0.017) |

For each procedure the FDR and the detection power (averaged over 25 runs) are displayed. Numbers in brackets correspond to standard errors. False discovery rate above 0.05 are in boldface.

**Table 16.** False discovery rate and detection power of the procedures in sparse and dense scenarios with *Q* = 2, *ρ* = 0.7 and *ξ* = 0.3 (settings 7s and 8s).

| Method | Sparse |  | Dense |  |
| --- | --- | --- | --- | --- |
|  | FDR | Power | FDR | Power |
| DACT_Efron | 0 (0) | 0 (0.001) | 0 (0) | 0 (0) |
| DACT_JC | 0 (0) | 0 (0.001) | 0.009 (0.029) | 0.001 (0.002) |
| DACT_NO | **0.249 (0.057)** | 0.176 (0.077) | **0.09 (0.009)** | 0.1 (0.02) |
| HDMT | **0.574 (0.027)** | 0.413 (0.048) | **0.459 (0.01)** | 0.634 (0.025) |
| PLACO | **0.655 (0.186)** | 0.009 (0.006) | 0.058 (0.021) | 0.01 (0.004) |
| adaFilter | **0.382 (0.034)** | 0.264 (0.052) | **0.146 (0.012)** | 0.208 (0.019) |
| qch | **0.553 (0.05)** | 0.402 (0.063) | **0.353 (0.023)** | 0.422 (0.032) |
| IMIX | **0.912 (0.007)** | 0.876 (0.042) | **0.351 (0.039)** | 0.583 (0.04) |
| c-csmGmm | 0.057 (0.059) | 0.006 (0.004) | 0.059 (0.01) | 0.048 (0.01) |
| Primo | **0.096 (0.044)** | 0.05 (0.019) | **0.144 (0.012)** | 0.19 (0.015) |
| qch_copula | **0.089 (0.086)** | 0.039 (0.025) | **0.065 (0.008)** | 0.124 (0.017) |

For each procedure the FDR and the detection power (averaged over 25 runs) are displayed. Numbers in brackets correspond to standard errors. False discovery rate above 0.05 are in boldface.

**Fig. 4.**
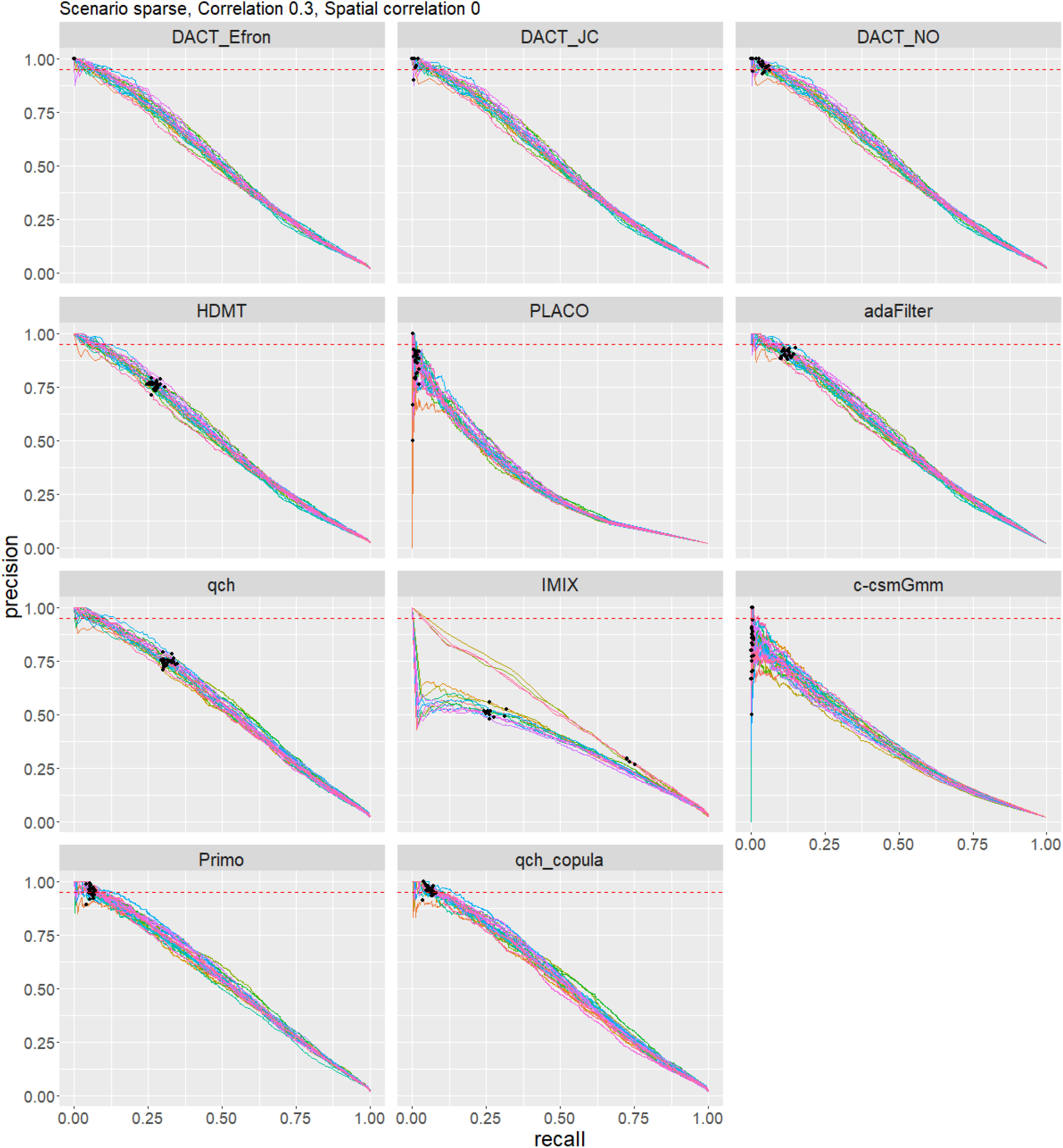
Precision-recall curves for simulation setting 3 (*Q* = 2, *ρ* = 0.3,*ξ* = 0,sparse), shown for each method. Each curve corresponds to one of the 25 simulation replicates. The red dotted line indicates the nominal precision level of 0.95. Black points indicates the threshold used by each method to control of the false discovery rate at the nominal level of 0.05.

**Fig. 5.**
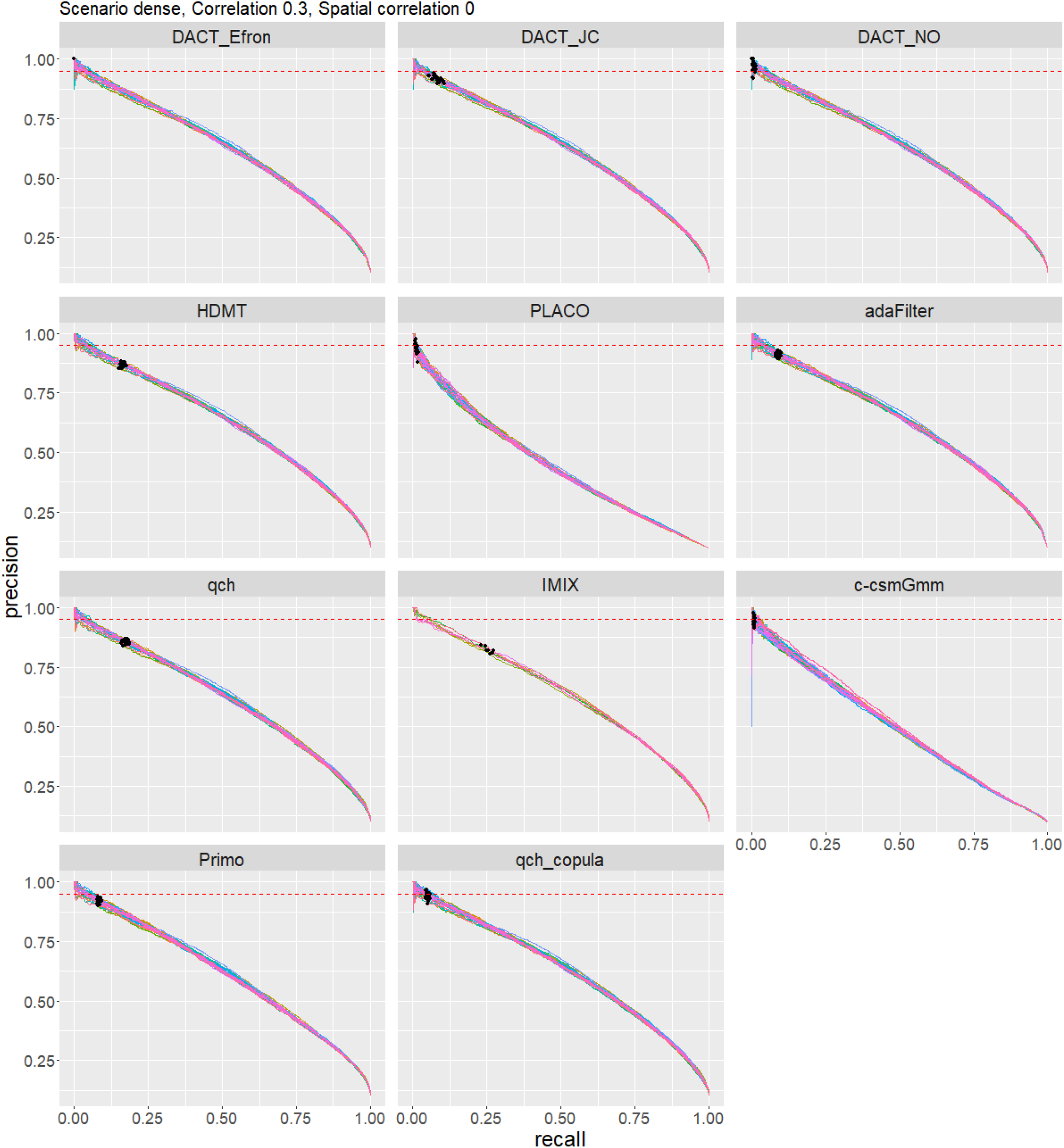
Precision-recall curves for simulation setting 4 (*Q* = 2, *ρ* = 0.3,*ξ* = 0, dense), shown for each method. Each curve corresponds to one of the 25 simulation replicates. The red dotted line indicates the nominal precision level of 0.95. Black points indicates the threshold used by each method to control of the false discovery rate at the nominal level of 0.05.

**Table 17.** Performance of the procedures Primo, adaFilter and qch_copula for the simulation settings 9-10 (*Q* = 8) and 17-18 (*Q* = 16) with *ρ* = 0 and *ξ* = 0.

| $Q$ | Scenario | Test | Primo | | adaFilter | | qch_copula | |
| --- | --- | --- | --- | --- | --- | --- | --- | --- |
|  |  |  | FDR | Power | FDR | Power | FDR | Power |
| 8 | Dense | at least 2 | 0.013 (0.011) | 0.083 (0.069) | 0.005 (0.001) | 0.317 (0.004) | 0.036 (0.005) | 0.77 (0.013) |
|  |  | at least 4 | 0.003 (0.003) | 0.042 (0.035) | 0 (0) | 0.025 (0.003) | 0.021 (0.004) | 0.776 (0.018) |
|  |  | at least 8 | 0.003 (0.005) | 0.01 (0.008) | 0 (0) | 0 (0) | 0.014 (0.005) | 0.124 (0.05) |
|  | Sparse | at least 2 | 0.011 (0.01) | 0.062 (0.052) | 0.009 (0.002) | 0.26 (0.008) | 0.039 (0.003) | 0.685 (0.006) |
|  |  | at least 4 | 0.002 (0.003) | 0.045 (0.038) | 0 (0) | 0.026 (0.008) | 0.038 (0.005) | 0.804 (0.015) |
|  |  | at least 8 | 0.003 (0.007) | 0.015 (0.014) | 0 (0) | 0 (0.001) | 0.03 (0.018) | 0.21 (0.062) |
| 16 | Dense | at least 2 | - | - | 0.004 (0.001) | 0.309 (0.003) | 0.041 (0.001) | 0.724 (0.003) |
|  |  | at least 4 | - | - | 0 (0) | 0.101 (0.004) | 0.038 (0.002) | 0.793 (0.005) |
|  |  | at least 8 | - | - | 0 (0) | 0.001 (0.001) | 0.037 (0.002) | 0.898 (0.005) |
|  |  | at least 14 | - | - | 0 (0) | 0 (0) | 0.003 (0.002) | 0.852 (0.043) |
|  |  | at least 16 | - | - | 0 (0) | 0 (0) | 0.02 (0.015) | 0.02 (0.013) |
|  | Sparse | at least 2 | - | - | 0.008 (0.002) | 0.264 (0.007) | 0.049 (0.003) | 0.653 (0.004) |
|  |  | at least 4 | - | - | 0 (0) | 0.101 (0.008) | 0.059 (0.003) | 0.812 (0.006) |
|  |  | at least 8 | - | - | 0 (0) | 0.002 (0.002) | 0.043 (0.004) | 0.941 (0.01) |
|  |  | at least 14 | - | - | 0 (0) | 0 (0) | 0.006 (0.004) | 0.883 (0.055) |
|  |  | at least 16 | - | - | 0 (0) | 0 (0) | 0.058 (0.052) | 0.118 (0.065) |

For each procedure the FDR and the detection power (averaged over 25 runs) are displayed. Numbers in brackets correspond to standard errors.

**Table 18.** Performance of the procedures Primo, adaFilter and qch_copula for the simulation settings 13-14 (*Q* = 8) and 21-22 (*Q* = 16) with *ρ* = 0.5 and *ξ* = 0.

| $Q$ | Scenario | Test | Primo | | adaFilter | | qch_copula | |
| --- | --- | --- | --- | --- | --- | --- | --- | --- |
|  |  |  | FDR | Power | FDR | Power | FDR | Power |
| 8 | Dense | at least 2 | 0.005 (0.001) | 0.144 (0.004) | 0.03 (0.002) | 0.315 (0.003) | 0.058 (0.002) | 0.617 (0.004) |
|  |  | at least 4 | 0.006 (0.001) | 0.147 (0.005) | 0.02 (0.002) | 0.199 (0.003) | **0.064 (0.003)** | 0.549 (0.005) |
|  |  | at least 8 | **0.075 (0.006)** | 0.178 (0.006) | 0.043 (0.005) | 0.111 (0.008) | **0.104 (0.007)** | 0.28 (0.006) |
|  | Sparse | at least 2 | 0.008 (0.003) | 0.115 (0.01) | **0.126 (0.008)** | 0.29 (0.008) | **0.063 (0.005)** | 0.422 (0.007) |
|  |  | at least 4 | 0.01 (0.004) | 0.12 (0.011) | **0.076 (0.008)** | 0.196 (0.006) | **0.067 (0.007)** | 0.415 (0.011) |
|  |  | at least 8 | **0.142 (0.031)** | 0.09 (0.01) | **0.078 (0.019)** | 0.107 (0.009) | **0.104 (0.023)** | 0.119 (0.012) |
| 16 | Dense | at least 2 | - | - | 0.034 (0.002) | 0.265 (0.003) | 0.057 (0.002) | 0.722 (0.004) |
|  |  | at least 4 | - | - | 0.025 (0.002) | 0.182 (0.003) | **0.067 (0.002)** | 0.756 (0.004) |
|  |  | at least 8 | - | - | 0.028 (0.003) | 0.139 (0.004) | **0.083 (0.002)** | 0.681 (0.005) |
|  |  | at least 14 | - | - | 0.009 (0.004) | 0.061 (0.004) | **0.086 (0.003)** | 0.528 (0.008) |
|  |  | at least 16 | - | - | **0.063 (0.013)** | 0.056 (0.007) | **0.124 (0.011)** | 0.23 (0.014) |
|  | Sparse | at least 2 | - | - | **0.153 (0.008)** | 0.246 (0.006) | **0.08 (0.007)** | 0.647 (0.007) |
|  |  | at least 4 | - | - | **0.105 (0.008)** | 0.18 (0.006) | **0.094 (0.008)** | 0.724 (0.009) |
|  |  | at least 8 | - | - | **0.072 (0.008)** | 0.138 (0.008) | **0.1 (0.008)** | 0.582 (0.014) |
|  |  | at least 14 | - | - | 0.033 (0.017) | 0.06 (0.009) | **0.086 (0.016)** | 0.336 (0.017) |
|  |  | at least 16 | - | - | **0.083 (0.036)** | 0.06 (0.012) | **0.172 (0.062)** | 0.096 (0.023) |

For each procedure the FDR and the detection power (averaged over 25 runs) are displayed. Numbers in brackets correspond to standard errors.

**Table 19.** Performance of the procedures Primo, adaFilter and qch_copula for the simulation settings 15-16 (*Q* = 8) and 23-24 (*Q* = 16) with *ρ* = 0.7 and *ξ* = 0.

| $Q$ | Scenario | Test | Primo | | adaFilter | | qch_copula | |
| --- | --- | --- | --- | --- | --- | --- | --- | --- |
|  |  |  | FDR | Power | FDR | Power | FDR | Power |
| 8 | Dense | at least 2 | 0.007 (0.006) | 0.084 (0.07) | 0.039 (0.002) | 0.312 (0.003) | 0.056 (0.006) | 0.755 (0.024) |
|  |  | at least 4 | 0.009 (0.008) | 0.096 (0.08) | 0.04 (0.002) | 0.244 (0.003) | **0.062 (0.013)** | 0.709 (0.021) |
|  |  | at least 8 | **0.061 (0.051)** | 0.128 (0.106) | **0.098 (0.006)** | 0.231 (0.007) | **0.118 (0.059)** | 0.305 (0.036) |
|  | Sparse | at least 2 | 0.008 (0.007) | 0.075 (0.063) | **0.19 (0.006)** | 0.295 (0.006) | **0.065 (0.007)** | 0.635 (0.011) |
|  |  | at least 4 | 0.011 (0.011) | 0.078 (0.066) | **0.171 (0.006)** | 0.239 (0.008) | **0.066 (0.012)** | 0.591 (0.017) |
|  |  | at least 8 | **0.136 (0.117)** | 0.046 (0.039) | **0.243 (0.02)** | 0.221 (0.011) | **0.116 (0.094)** | 0.057 (0.041) |
| 16 | Dense | at least 2 | - | - | 0.041 (0.002) | 0.252 (0.002) | **0.171 (0.222)** | 0.761 (0.204) |
|  |  | at least 4 | - | - | 0.044 (0.002) | 0.198 (0.003) | 0.05 (0.003) | 0.892 (0.004) |
|  |  | at least 8 | - | - | **0.064 (0.006)** | 0.177 (0.004) | 0.053 (0.002) | 0.777 (0.005) |
|  |  | at least 14 | - | - | 0.047 (0.005) | 0.137 (0.005) | 0.043 (0.003) | 0.6 (0.008) |
|  |  | at least 16 | - | - | **0.113 (0.009)** | 0.168 (0.007) | 0.046 (0.007) | 0.179 (0.011) |
|  | Sparse | at least 2 | - | - | **0.209 (0.009)** | 0.243 (0.006) | **0.242 (0.329)** | 0.65 (0.281) |
|  |  | at least 4 | - | - | **0.201 (0.01)** | 0.196 (0.005) | 0.058 (0.015) | 0.808 (0.007) |
|  |  | at least 8 | - | - | **0.204 (0.02)** | 0.175 (0.006) | 0.047 (0.013) | 0.617 (0.013) |
|  |  | at least 14 | - | - | **0.176 (0.021)** | 0.131 (0.008) | 0.029 (0.01) | 0.391 (0.023) |
|  |  | at least 16 | - | - | **0.286 (0.028)** | 0.156 (0.01) | 0 (0) | 0 (0) |

For each procedure the FDR and the detection power (averaged over 25 runs) are displayed. Numbers in brackets correspond to standard errors.

**Table 20.** Performance of the procedures Primo, adaFilter and qch_copula for the simulation settings 9s-10s (*Q* = 8) and 17s-18s (*Q* = 16) with *ρ* = 0 and *ξ* = 0.3.

| $Q$ | Scenario | Test | Primo | | adaFilter | | qch_copula | |
| --- | --- | --- | --- | --- | --- | --- | --- | --- |
|  |  |  | FDR | Power | FDR | Power | FDR | Power |
| 8 | Dense | at least 2 | 0.011 (0.01) | 0.083 (0.07) | 0.005 (0.001) | 0.317 (0.011) | 0.034 (0.008) | 0.75 (0.034) |
|  |  | at least 4 | 0.003 (0.003) | 0.043 (0.036) | 0 (0) | 0.025 (0.006) | 0.016 (0.009) | 0.715 (0.075) |
|  |  | at least 8 | 0.001 (0.003) | 0.008 (0.007) | 0 (0) | 0 (0) | 0.011 (0.013) | 0.093 (0.097) |
|  | Sparse | at least 2 | 0.01 (0.009) | 0.065 (0.055) | 0.009 (0.002) | 0.264 (0.025) | 0.038 (0.009) | 0.683 (0.021) |
|  |  | at least 4 | 0.002 (0.003) | 0.045 (0.04) | 0 (0.001) | 0.029 (0.012) | 0.034 (0.011) | 0.78 (0.036) |
|  |  | at least 8 | 0.005 (0.018) | 0.013 (0.013) | 0 (0) | 0 (0) | 0.041 (0.033) | 0.178 (0.121) |
| 16 | Dense | at least 2 | - | - | 0.02 (0.002) | 0.276 (0.011) | 0.054 (0.004) | 0.667 (0.015) |
|  |  | at least 4 | - | - | 0.007 (0.002) | 0.164 (0.009) | 0.053 (0.005) | 0.676 (0.02) |
|  |  | at least 8 | - | - | 0.006 (0.003) | 0.09 (0.009) | 0.05 (0.005) | 0.584 (0.031) |
|  |  | at least 14 | - | - | 0.001 (0.002) | 0.014 (0.004) | 0.019 (0.002) | 0.354 (0.042) |
|  |  | at least 16 | - | - | 0.027 (0.037) | 0.008 (0.005) | 0.042 (0.021) | 0.061 (0.038) |
|  | Sparse | at least 2 | - | - | **0.074 (0.008)** | 0.25 (0.021) | **0.076 (0.009)** | 0.519 (0.031) |
|  |  | at least 4 | - | - | 0.023 (0.005) | 0.168 (0.018) | **0.08 (0.011)** | 0.575 (0.045) |
|  |  | at least 8 | - | - | 0.005 (0.005) | 0.095 (0.025) | **0.071 (0.013)** | 0.42 (0.06) |
|  |  | at least 14 | - | - | 0 (0) | 0.014 (0.009) | 0.02 (0.01) | 0.168 (0.061) |
|  |  | at least 16 | - | - | 0.002 (0.008) | 0.008 (0.007) | 0.056 (0.06) | 0.048 (0.052) |

For each procedure the FDR and the detection power (averaged over 25 runs) are displayed. Numbers in brackets correspond to standard errors.

**Table 21.** Performance of the procedures Primo, adaFilter and qch_copula for the simulation settings 13s-14s (*Q* = 8) and 21s-22s (*Q* = 16) with *ρ* = 0.5 and *ξ* = 0.3.

| $Q$ | Scenario | Test | Primo | | adaFilter | | qch_copula | |
| --- | --- | --- | --- | --- | --- | --- | --- | --- |
|  |  |  | FDR | Power | FDR | Power | FDR | Power |
| 8 | Dense | at least 2 | 0.003 (0.003) | 0.087 (0.072) | 0.029 (0.003) | 0.313 (0.01) | 0.048 (0.006) | 0.614 (0.019) |
|  |  | at least 4 | 0.004 (0.004) | 0.087 (0.073) | 0.02 (0.003) | 0.196 (0.01) | 0.042 (0.006) | 0.528 (0.019) |
|  |  | at least 8 | 0.045 (0.038) | 0.105 (0.088) | 0.041 (0.009) | 0.112 (0.015) | 0.044 (0.007) | 0.272 (0.058) |
|  | Sparse | at least 2 | 0.004 (0.004) | 0.073 (0.062) | **0.126 (0.009)** | 0.288 (0.024) | 0.059 (0.013) | 0.432 (0.031) |
|  |  | at least 4 | 0.006 (0.006) | 0.073 (0.062) | **0.074 (0.009)** | 0.195 (0.023) | 0.054 (0.016) | 0.412 (0.056) |
|  |  | at least 8 | **0.087 (0.077)** | 0.053 (0.048) | **0.075 (0.019)** | 0.106 (0.026) | **0.06 (0.035)** | 0.13 (0.066) |
| 16 | Dense | at least 2 | - | - | 0.035 (0.003) | 0.263 (0.007) | 0.055 (0.006) | 0.744 (0.017) |
|  |  | at least 4 | - | - | 0.026 (0.004) | 0.181 (0.007) | 0.052 (0.007) | 0.752 (0.021) |
|  |  | at least 8 | - | - | 0.03 (0.007) | 0.137 (0.011) | 0.054 (0.005) | 0.606 (0.024) |
|  |  | at least 14 | - | - | 0.01 (0.005) | 0.06 (0.008) | 0.031 (0.006) | 0.341 (0.038) |
|  |  | at least 16 | - | - | 0.058 (0.02) | 0.058 (0.015) | 0.041 (0.01) | 0.123 (0.038) |
|  | Sparse | at least 2 | - | - | **0.149 (0.012)** | 0.245 (0.015) | **0.085 (0.024)** | 0.631 (0.031) |
|  |  | at least 4 | - | - | **0.106 (0.011)** | 0.178 (0.016) | **0.089 (0.03)** | 0.663 (0.049) |
|  |  | at least 8 | - | - | **0.071 (0.018)** | 0.141 (0.022) | **0.071 (0.02)** | 0.413 (0.045) |
|  |  | at least 14 | - | - | 0.027 (0.019) | 0.065 (0.02) | 0.023 (0.015) | 0.128 (0.048) |
|  |  | at least 16 | - | - | **0.082 (0.051)** | 0.064 (0.036) | 0.025 (0.055) | 0.019 (0.035) |

For each procedure the FDR and the detection power (averaged over 25 runs) are displayed. Numbers in brackets correspond to standard errors.

**Table 22.**
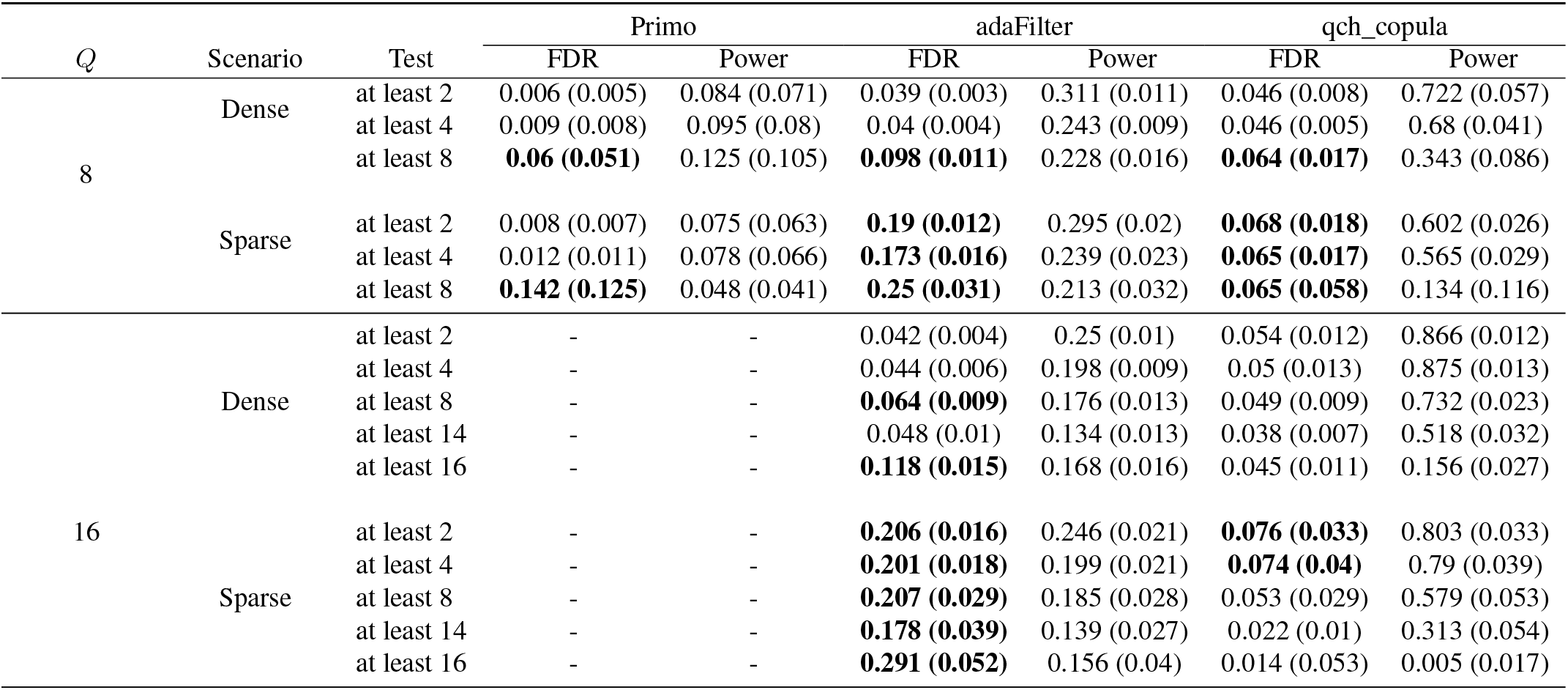
Performance of the procedures Primo, adaFilter and qch_copula for the simulation settings 15s-16s (*Q* = 8) and 23s-24s (*Q* = 16) with *ρ* = 0.7 and *ξ* = 0.3.

| $Q$ | Scenario | Test | Primo | | adaFilter | | qch_copula | |
| --- | --- | --- | --- | --- | --- | --- | --- | --- |
|  |  |  | FDR | Power | FDR | Power | FDR | Power |
| 8 | Dense | at least 2 | 0.006 (0.005) | 0.084 (0.071) | 0.039 (0.003) | 0.311 (0.011) | 0.046 (0.008) | 0.722 (0.057) |
|  |  | at least 4 | 0.009 (0.008) | 0.095 (0.08) | 0.04 (0.004) | 0.243 (0.009) | 0.046 (0.005) | 0.68 (0.041) |
|  |  | at least 8 | **0.06 (0.051)** | 0.125 (0.105) | **0.098 (0.011)** | 0.228 (0.016) | **0.064 (0.017)** | 0.343 (0.086) |
|  | Sparse | at least 2 | 0.008 (0.007) | 0.075 (0.063) | **0.19 (0.012)** | 0.295 (0.02) | **0.068 (0.018)** | 0.602 (0.026) |
|  |  | at least 4 | 0.012 (0.011) | 0.078 (0.066) | **0.173 (0.016)** | 0.239 (0.023) | **0.065 (0.017)** | 0.565 (0.029) |
|  |  | at least 8 | **0.142 (0.125)** | 0.048 (0.041) | **0.25 (0.031)** | 0.213 (0.032) | **0.065 (0.058)** | 0.134 (0.116) |
| 16 | Dense | at least 2 | - | - | 0.042 (0.004) | 0.25 (0.01) | 0.054 (0.012) | 0.866 (0.012) |
|  |  | at least 4 | - | - | 0.044 (0.006) | 0.198 (0.009) | 0.05 (0.013) | 0.875 (0.013) |
|  |  | at least 8 | - | - | **0.064 (0.009)** | 0.176 (0.013) | 0.049 (0.009) | 0.732 (0.023) |
|  |  | at least 14 | - | - | 0.048 (0.01) | 0.134 (0.013) | 0.038 (0.007) | 0.518 (0.032) |
|  |  | at least 16 | - | - | **0.118 (0.015)** | 0.168 (0.016) | 0.045 (0.011) | 0.156 (0.027) |
|  | Sparse | at least 2 | - | - | **0.206 (0.016)** | 0.246 (0.021) | **0.076 (0.033)** | 0.803 (0.033) |
|  |  | at least 4 | - | - | **0.201 (0.018)** | 0.199 (0.021) | **0.074 (0.04)** | 0.79 (0.039) |
|  |  | at least 8 | - | - | **0.207 (0.029)** | 0.185 (0.028) | 0.053 (0.029) | 0.579 (0.053) |
|  |  | at least 14 | - | - | **0.178 (0.039)** | 0.139 (0.027) | 0.022 (0.01) | 0.313 (0.054) |
|  |  | at least 16 | - | - | **0.291 (0.052)** | 0.156 (0.04) | 0.014 (0.053) | 0.005 (0.017) |

For each procedure the FDR and the detection power (averaged over 25 runs) are displayed. Numbers in brackets correspond to standard errors.

**Fig. 6.**
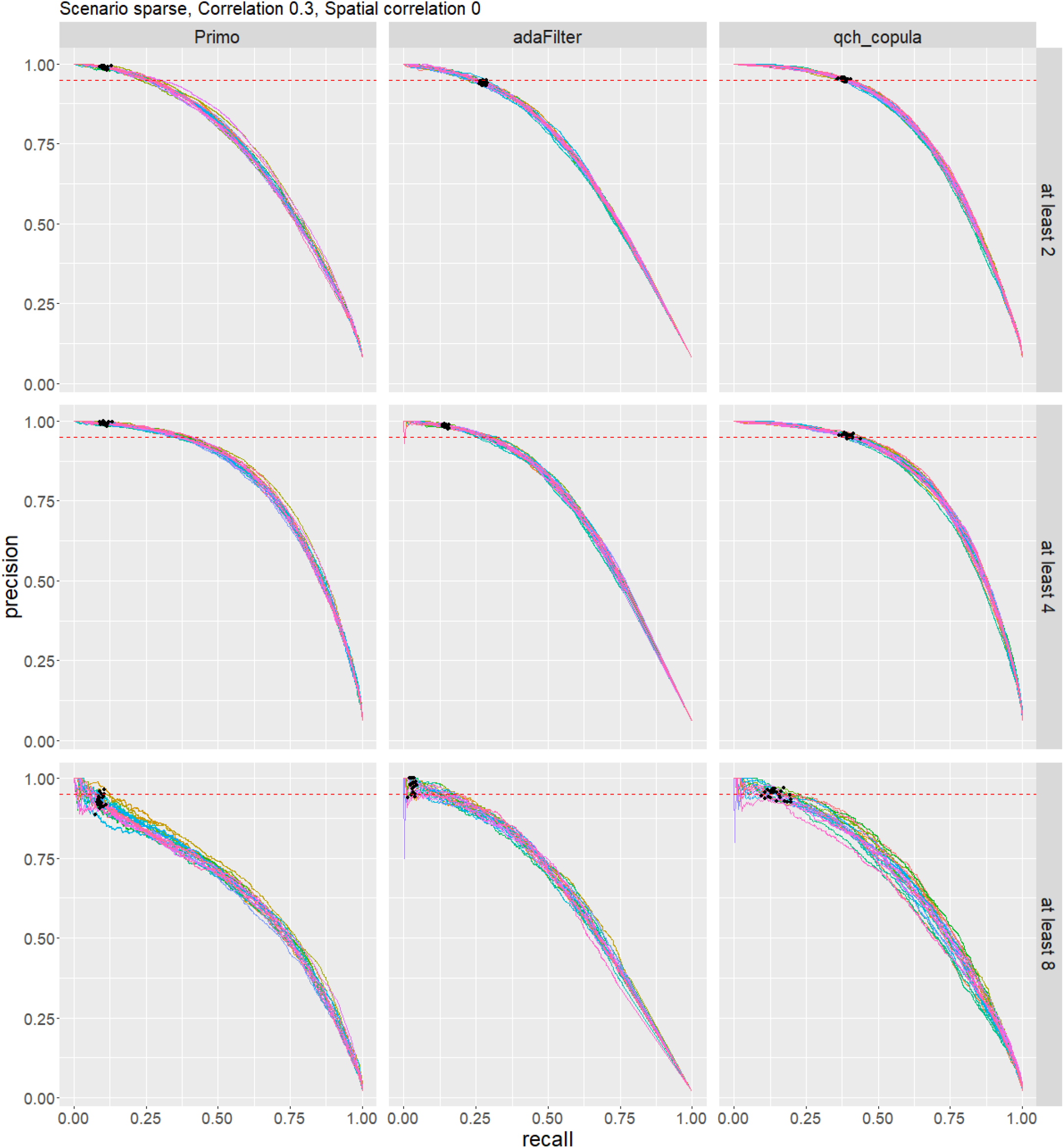
Precision-recall curves for simulation setting 11 (*Q* = 8, *ρ* = 0.3, *ξ* = 0, sparse), shown for each method. Each curve corresponds to one of the 25 simulation replicates. The red dotted line indicates the nominal precision level of 0.95. Black points indicates the threshold used by each method to control of the false discovery rate at the nominal level of 0.05.

**Fig. 7.**
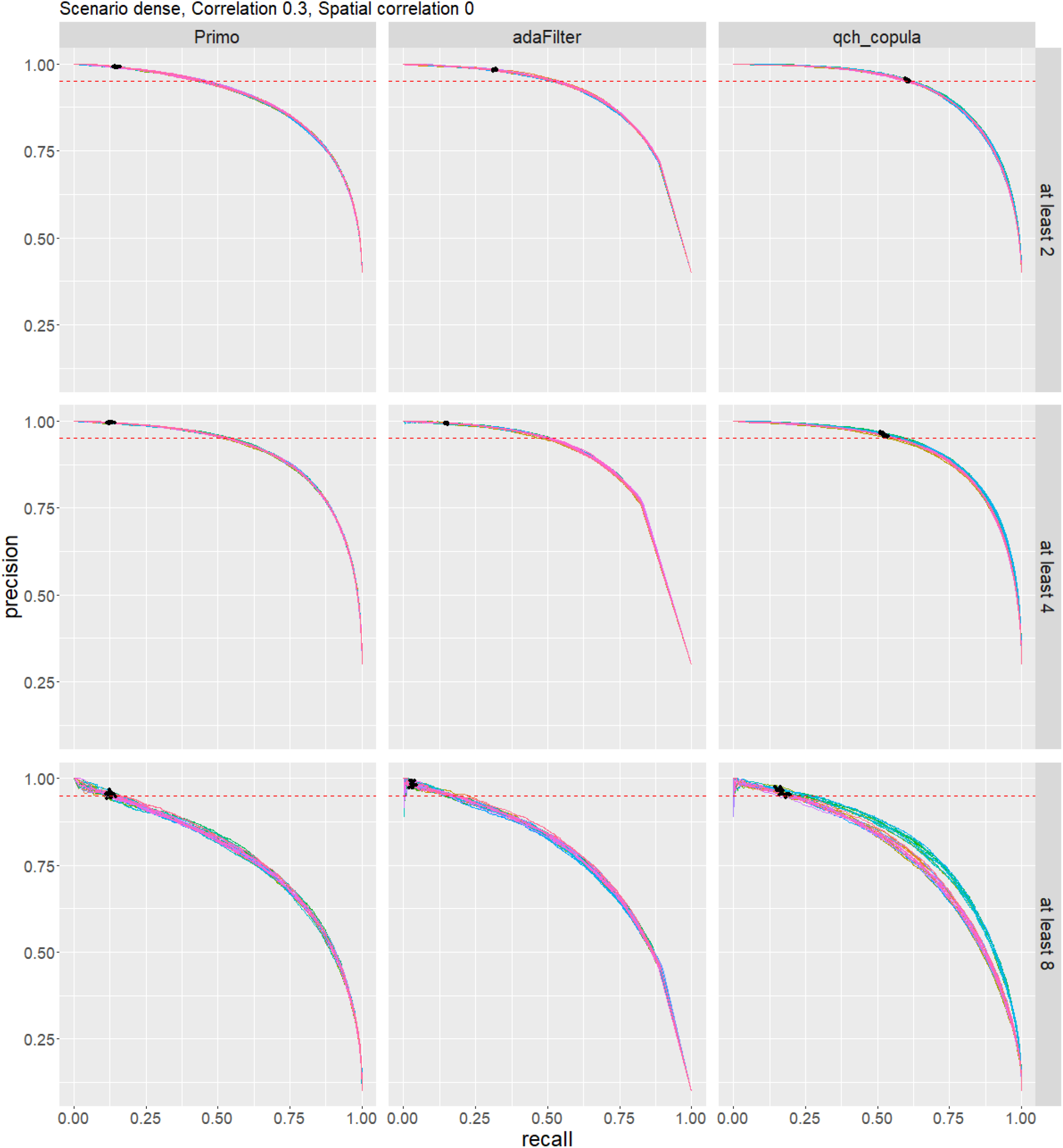
Precision-recall curves for simulation setting 12 (*Q* = 8, *ρ* = 0.3, *ξ* = 0, dense), shown for each method. Each curve corresponds to one of the 25 simulation replicates. The red dotted line indicates the nominal precision level of 0.95. Black points indicates the threshold used by each method to control of the false discovery rate at the nominal level of 0.05.

**Fig. 8.**
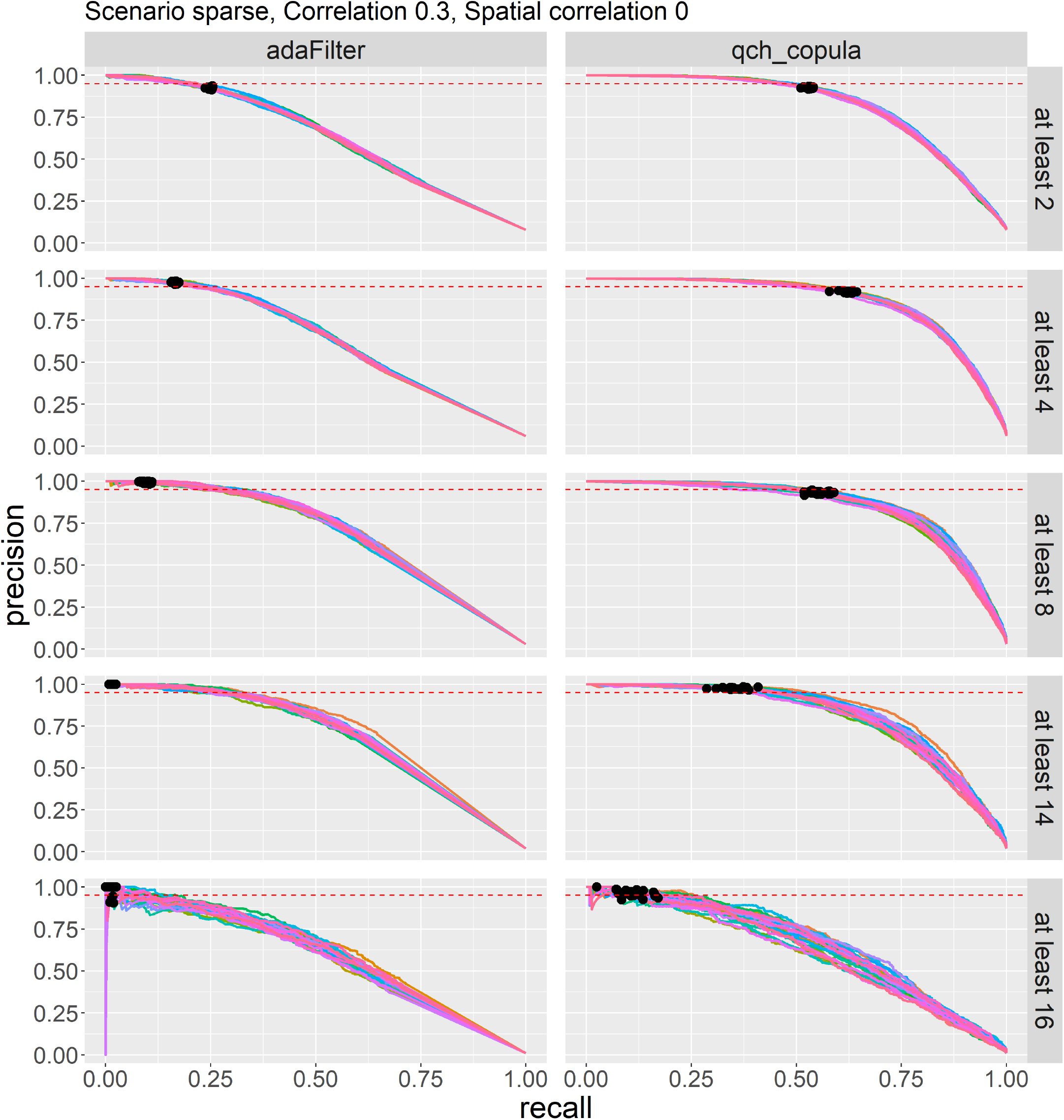
Precision-recall curves for simulation setting 19 (*Q* = 16, *ρ* = 0.3, *ξ* = 0, sparse), shown for each method. Each curve corresponds to one of the 25 simulation replicates. The red dotted line indicates the nominal precision level of 0.95. Black points indicates the threshold used by each method to control of the false discovery rate at the nominal level of 0.05.

**Fig. 9.**
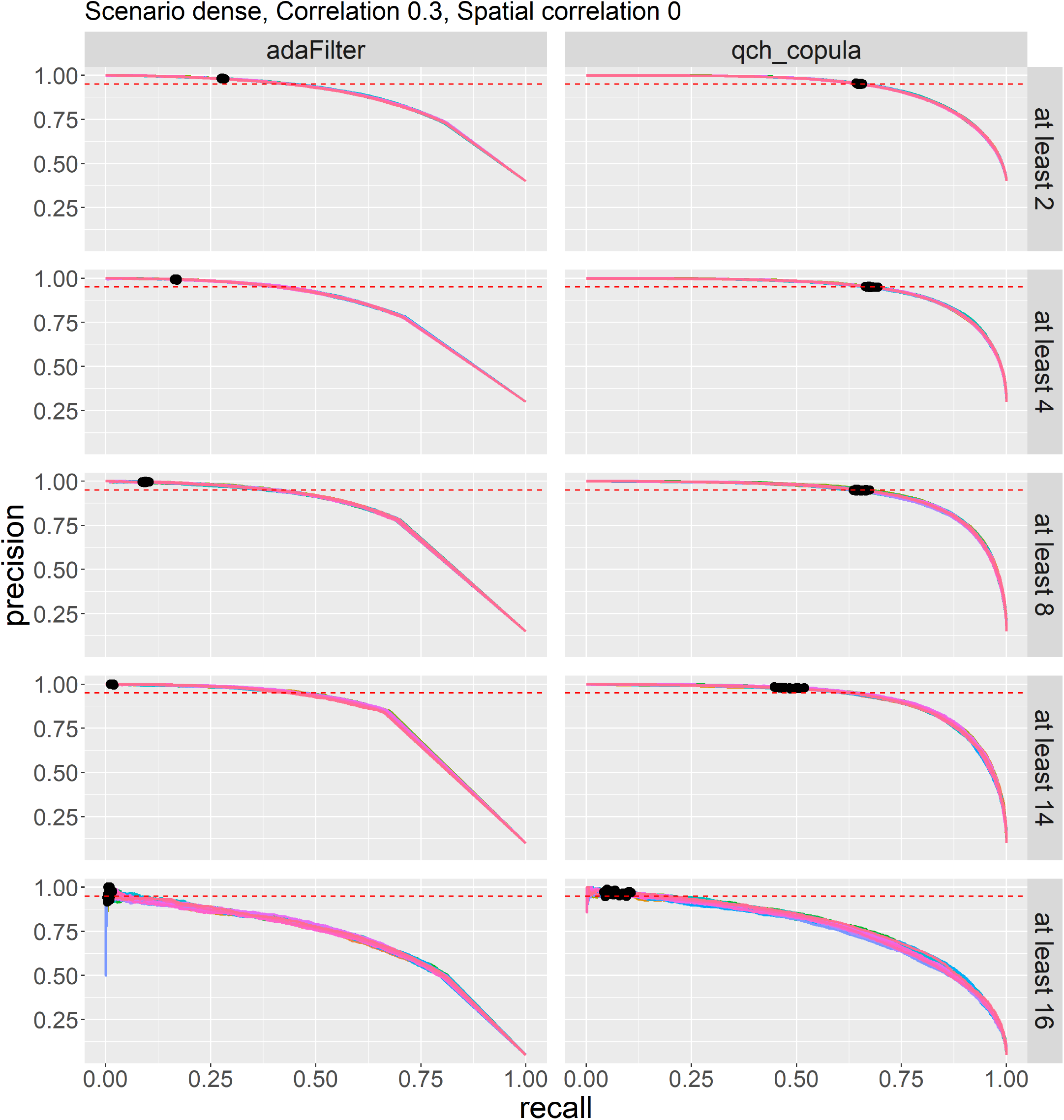
Precision-recall curves for simulation setting 20 (*Q* = 16, *ρ* = 0.3, *ξ* = 0, dense), shown for each method. Each curve corresponds to one of the 25 simulation replicates. The red dotted line indicates the nominal precision level of 0.95. Black points indicates the threshold used by each method to control of the false discovery rate at the nominal level of 0.05.

**Fig. 10.**
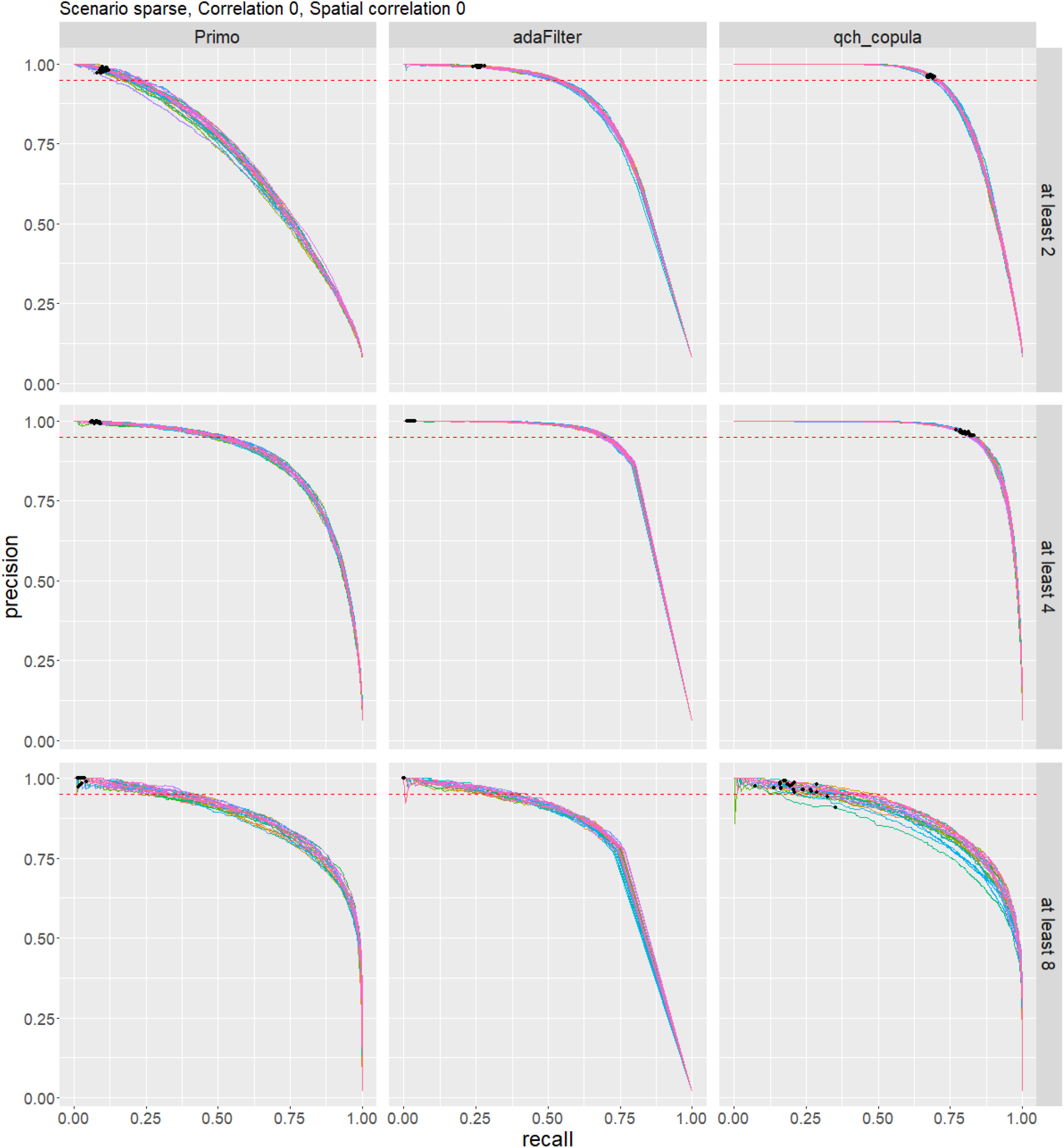
Precision-recall curves for simulation setting 9 (*Q* = 8, *ρ* = 0, *ξ* = 0, sparse), shown for each method. Each curve corresponds to one of the 25 simulation replicates. The red dotted line indicates the nominal precision level of 0.95. Black points indicates the threshold used by each method to control of the false discovery rate at the nominal level of 0.05.

**Fig. 11.**
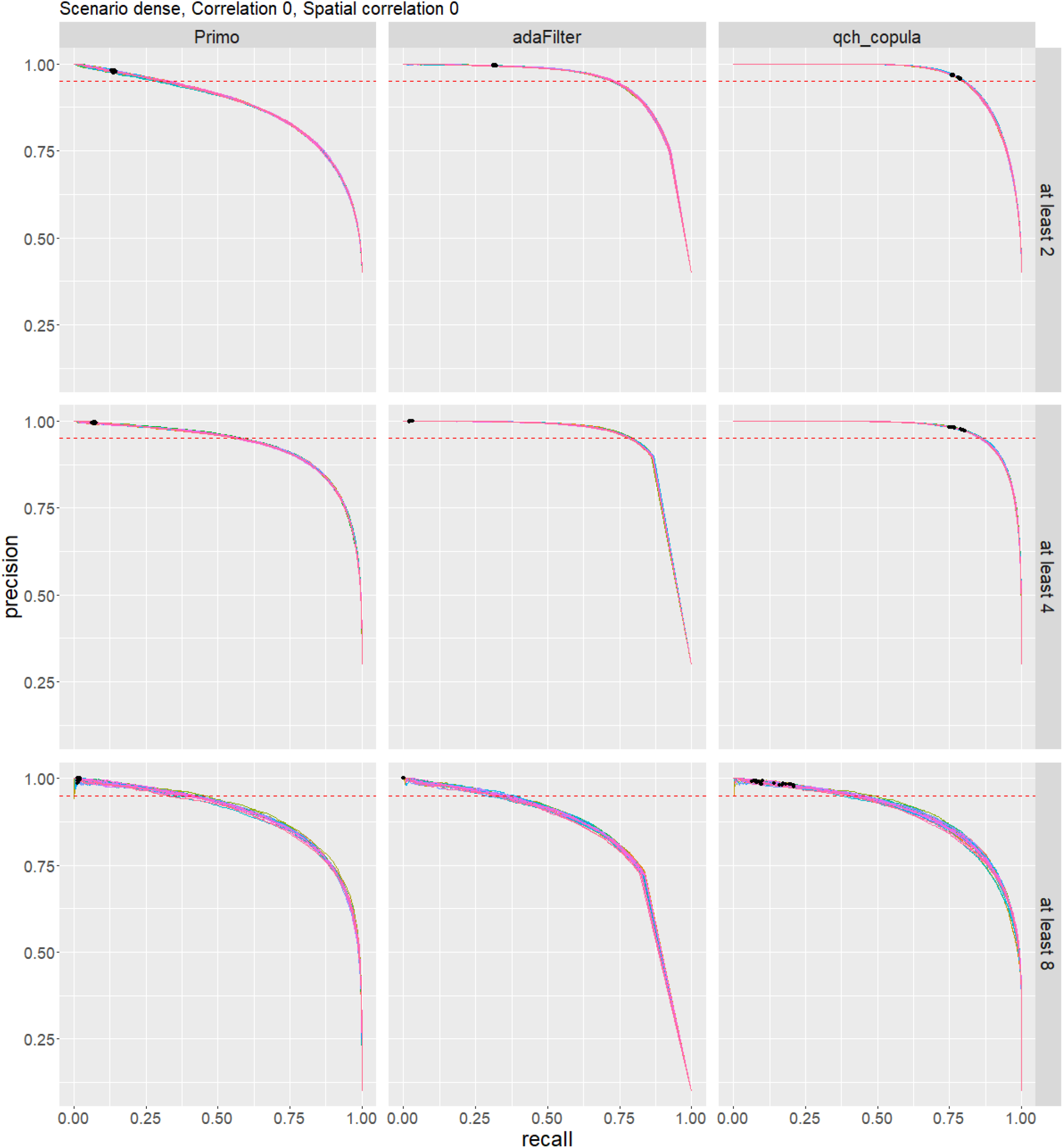
Precision-recall curves for simulation setting 10 (*Q* = 8, *ρ* = 0, *ξ* = 0, dense), shown for each method. Each curve corresponds to one of the 25 simulation replicates. The red dotted line indicates the nominal precision level of 0.95. Black points indicates the threshold used by each method to control of the false discovery rate at the nominal level of 0.05.

**Fig. 12.**
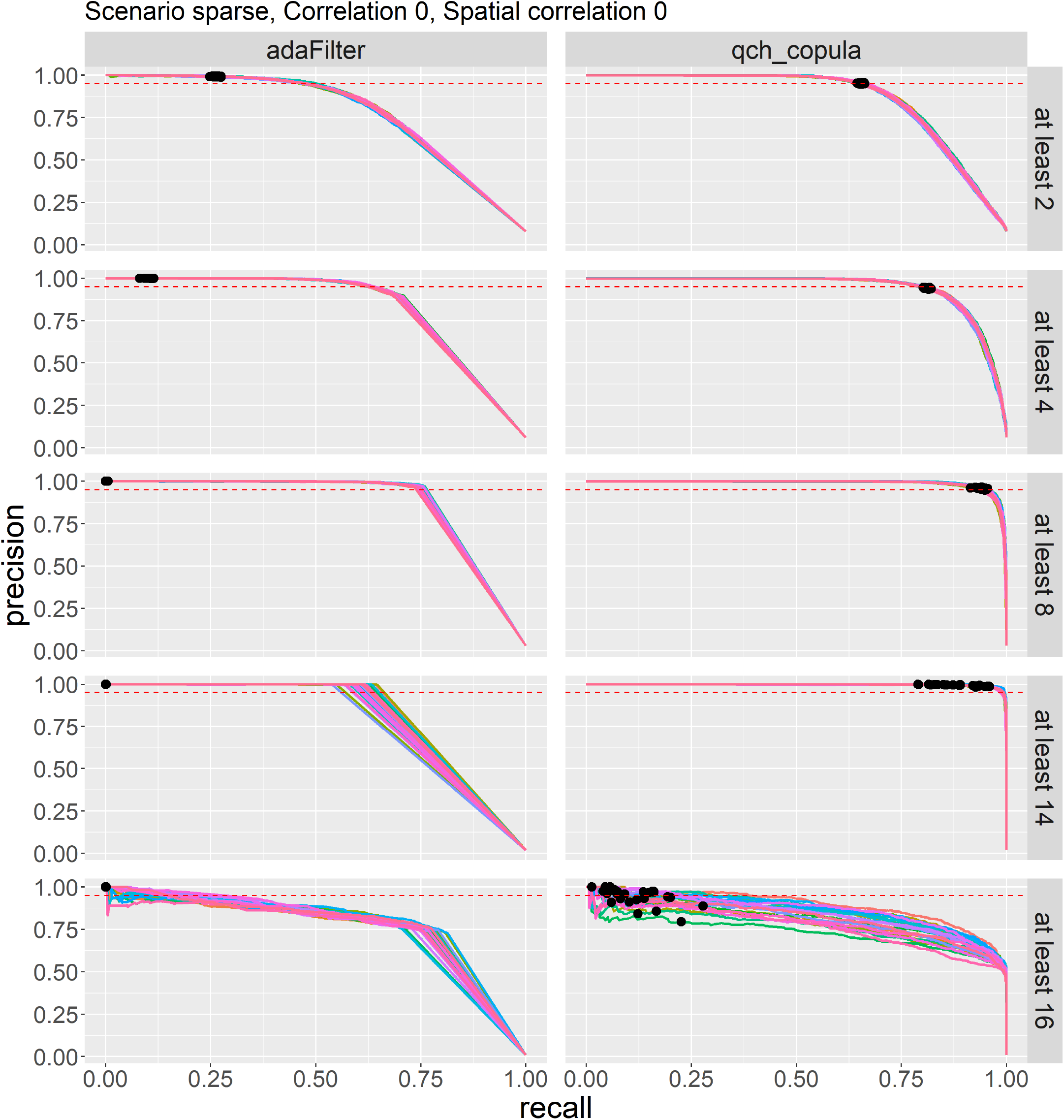
Precision-recall curves for simulation setting 17 (*Q* = 16, *ρ* = 0, *ξ* = 0, sparse), shown for each method. Each curve corresponds to one of the 25 simulation replicates. The red dotted line indicates the nominal precision level of 0.95. Black points indicates the threshold used by each method to control of the false discovery rate at the nominal level of 0.05.

**Fig. 13.**
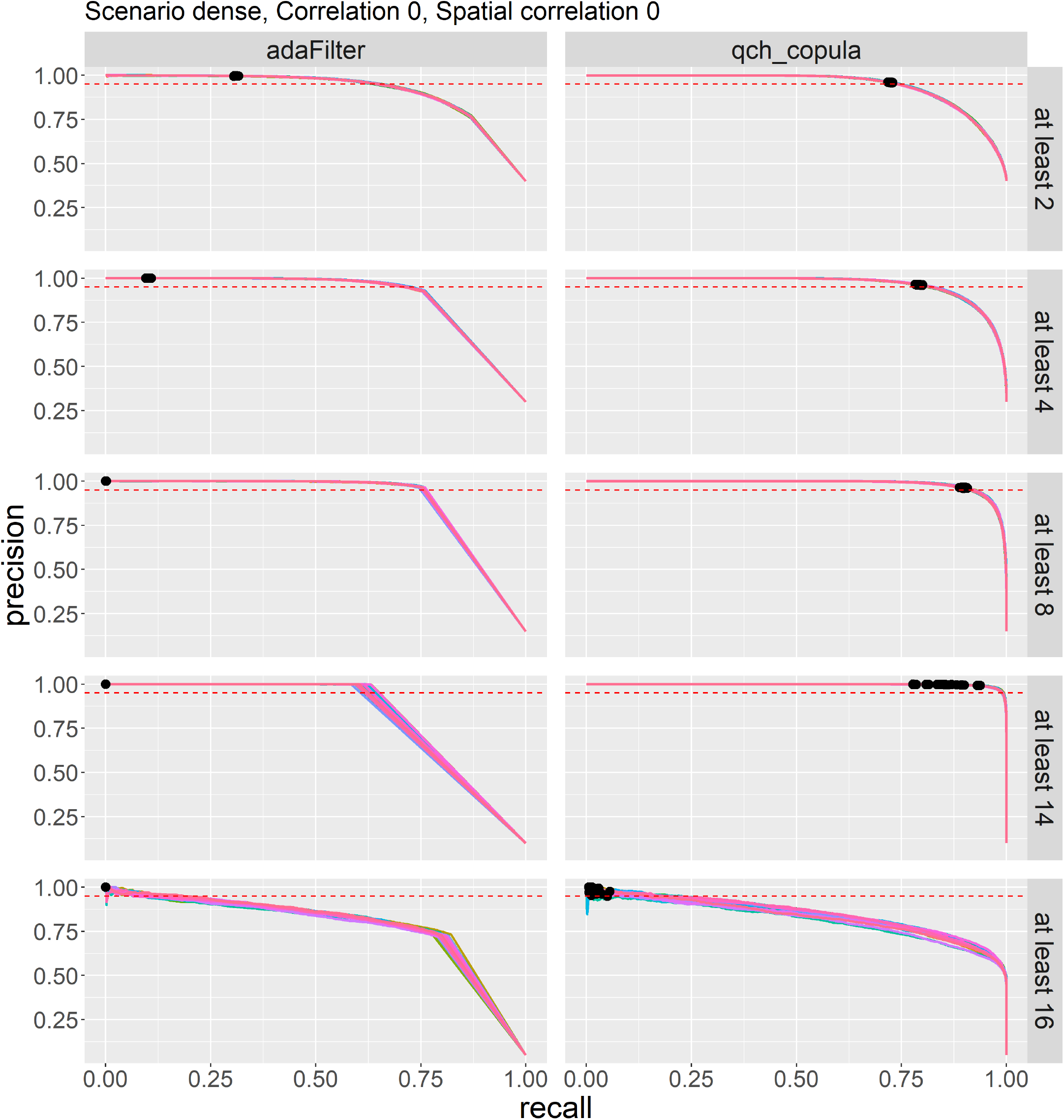
Precision-recall curves for simulation setting 16 (*Q* = 16, *ρ* = 0, *ξ* = 0, dense), shown for each method. Each curve corresponds to one of the 25 simulation replicates. The red dotted line indicates the nominal precision level of 0.95. Black points indicates the threshold used by each method to control of the false discovery rate at the nominal level of 0.05.

## Supplementary Note 6: Evaluation of marginal null proportion estimates

We compared the estimated marginal null proportions *π*_*q*_ to their true values across 25 simulations in both the sparse and dense scenarios (Fig. 14) for *Q* = 8. The visual inspection suggests a close alignment between the estimated and true values in both settings. To quantify estimation accuracy, we computed the mean squared error (MSE) for each simulation. In the sparse scenario, the average MSE was 2.18 *×* 10^−4^, and in the dense scenario it was at 6.94 *×* 10^−4^, indicating reliable estimation performance of the *π*_*q*_ parameters.

**Fig. 14.**
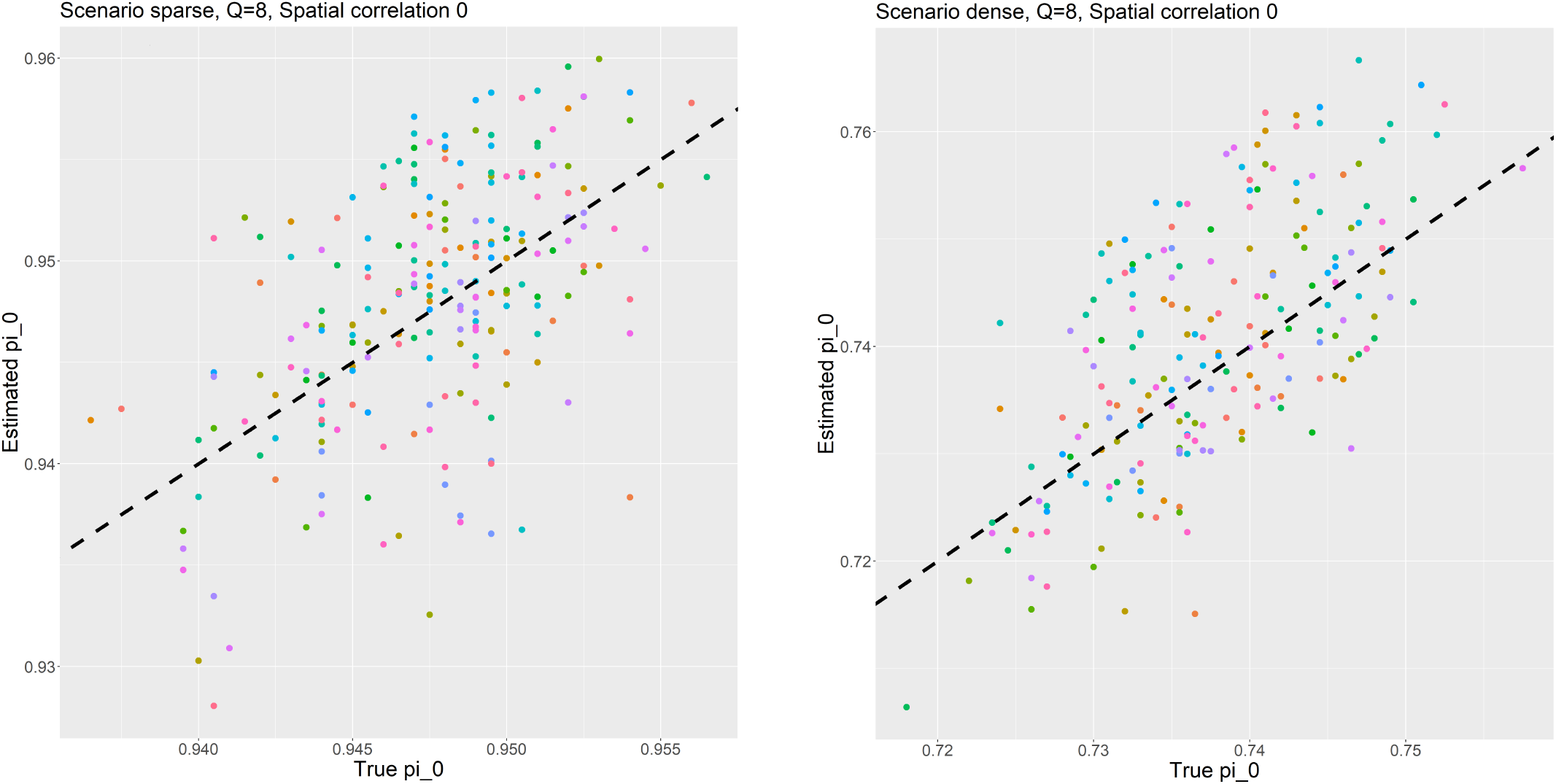
Estimated vs. true marginal null proportions *πq* across 25 simulation replicates under sparse and dense scenarios. Each color corresponds to a single simulation replicate. The black dotted line denotes the identity line (*y* = *x*), indicating perfect agreement between estimated and true values.

## Supplementary Note 7: Information on the datasets used for the application cases studies

### A. Application I: Detection of pleiotropic regions associated to psychiatric disorders

We analyzed 14 psychiatric disorders obtained from the Psychiatric Genomics Consortium (PGC) ^1^. The disorders studied are anorexia nervosa (AN), anxiety disorder (AD), attention-deficit/hyperactivity disorder (ADHD), autism spectrum disorder (ASD), alcohol use disorders (AUDIT-T alcohol use traits identification test based on total score, AUDIT-C alcohol use traits identification test based on consumption, AUDIT-P alcohol use traits identification test based on problematic consequences of drinking), bipolar disorder (BIP), cannabis use (CU^2^), major depression disorder (MDD), obsessive-compulsive disorder (OCD), post-traumatic stress disorder (PTSD), schizophrenia (SCZ) and Tourette’s syndrome (TS). Table 23 details the information about each study. For all the disorders, we obtained their European-only summary statistics and performed the following quality control:

i. excluded non-biallelic SNPs and those with strand-ambiguous alleles;
ii. removed duplicated SNPs.

**Table 23.** Summary information of 14 psychiatric disorders GWAS analyzed in this study.

| Traits | N(case/control) | PGC study | Reference |
| --- | --- | --- | --- |
| AN | 14,477 (3,495/10,982) | an2017 | (41) |
| AD | 17,526 (5,761/11,765) | anx2016 | (42) |
| ADHD | 53,293 (19,099/34,194) | adhd2019 | (43) |
| ASD | 46,350 (18,381/27,969) | asd2019 | (44) |
| AUDIT-T | 121,604 | sud2019-alcuse | (45) |
| AUDIT-C | 121,604 | sud2019-alcuse | (45) |
| AUDIT-P | 121,604 | sud2019-alcuse | (45) |
| BIP | 51,710 (20,352/31,358) | bip2019 | (46) |
| CU | 184,765 (53,180/131,585) | - | (47) |
| MDD | 480,359 (135,458/344,901) | mdd2018 | (48) |
| OCD | 9,725 (2,688/7,037) | ocd2018 | (49) |
| PTSD | 174,659 (23,212/151,447) | ptsd2019 | (50) |
| SCZ | 77,096 (33,640/43,456) | scz2019asi | (51) |
| TS | 14,307 (4,819/9,488) | ts2019 | (52) |

### B. Application II: Detection of viruses resistance hotspots regions in cucumber

Our second application case is based on GWAS summary statistics derived from the experiment described in (27). In this study, a panel of 226 elite cucumber lines from Bayer germplasm collections, 40 landraces from Bayer’s internal collections, and 23 hybrids-including resistance controls-were inoculated with six viruses to evaluate their responses. The viruses included zucchini yellow mosaic virus (ZYMV), papaya ringspot virus (PRSV), watermelon mosaic virus (WMV), cucumber mosaic virus (CMV), cucumber vein yellowing virus (CVYV), and cucumber green mottle mosaic virus (CGMMV). An individual GWAS was conducted on a number of SNPs ranging from *n* = 378, 049 to *n* = 424, 393 depending on the virus. The experimental protocol and results are fully detailed in (27). The aim of the study was to identify QTLs associated with virus resistance in cucumber.

## Supplementary Note 8: Supplementary figures for applications

**Fig. 15.**
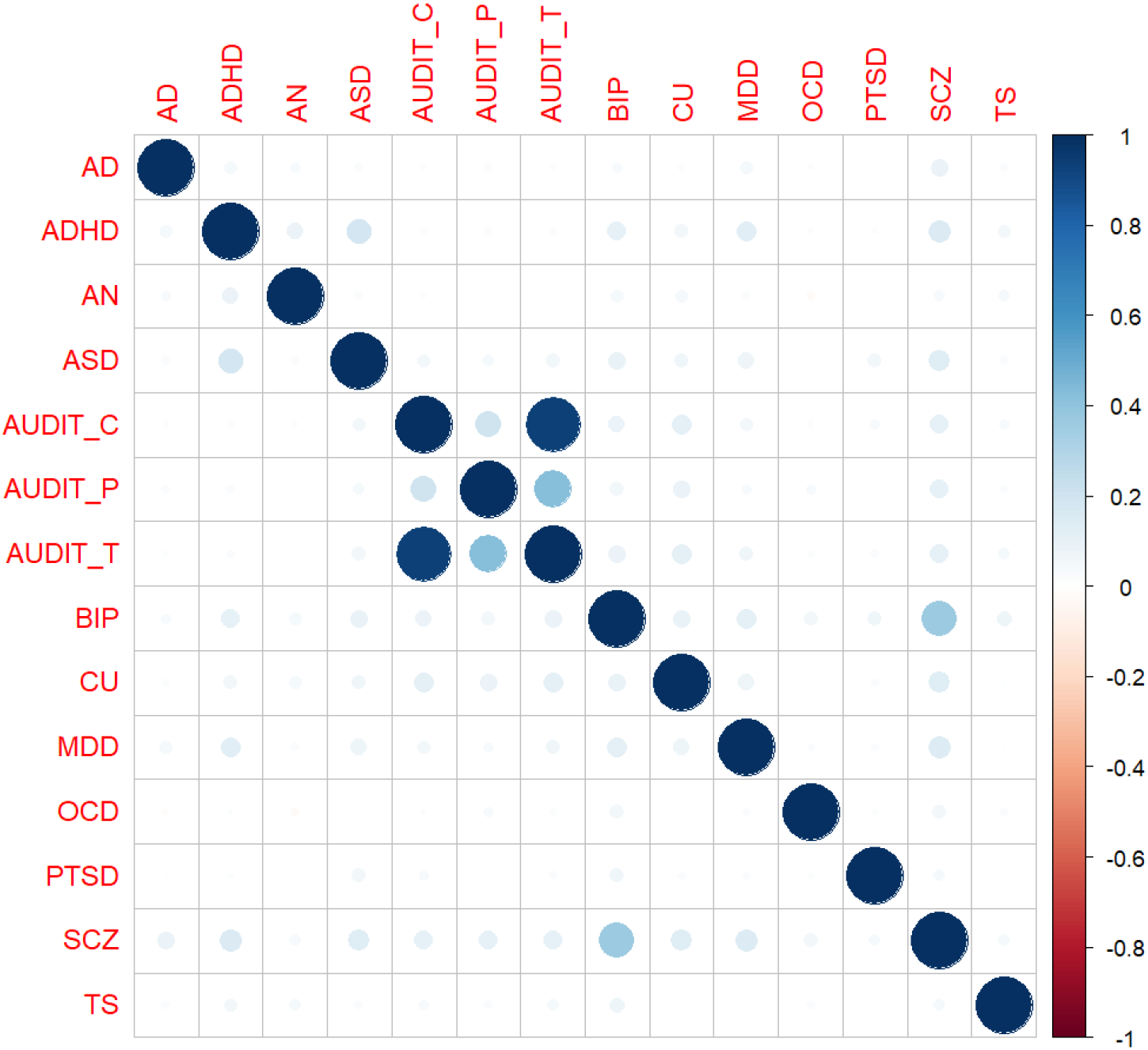
Estimated correlation matrix between the 14 psychiatric disorders GWAS.

**Fig. 16.**
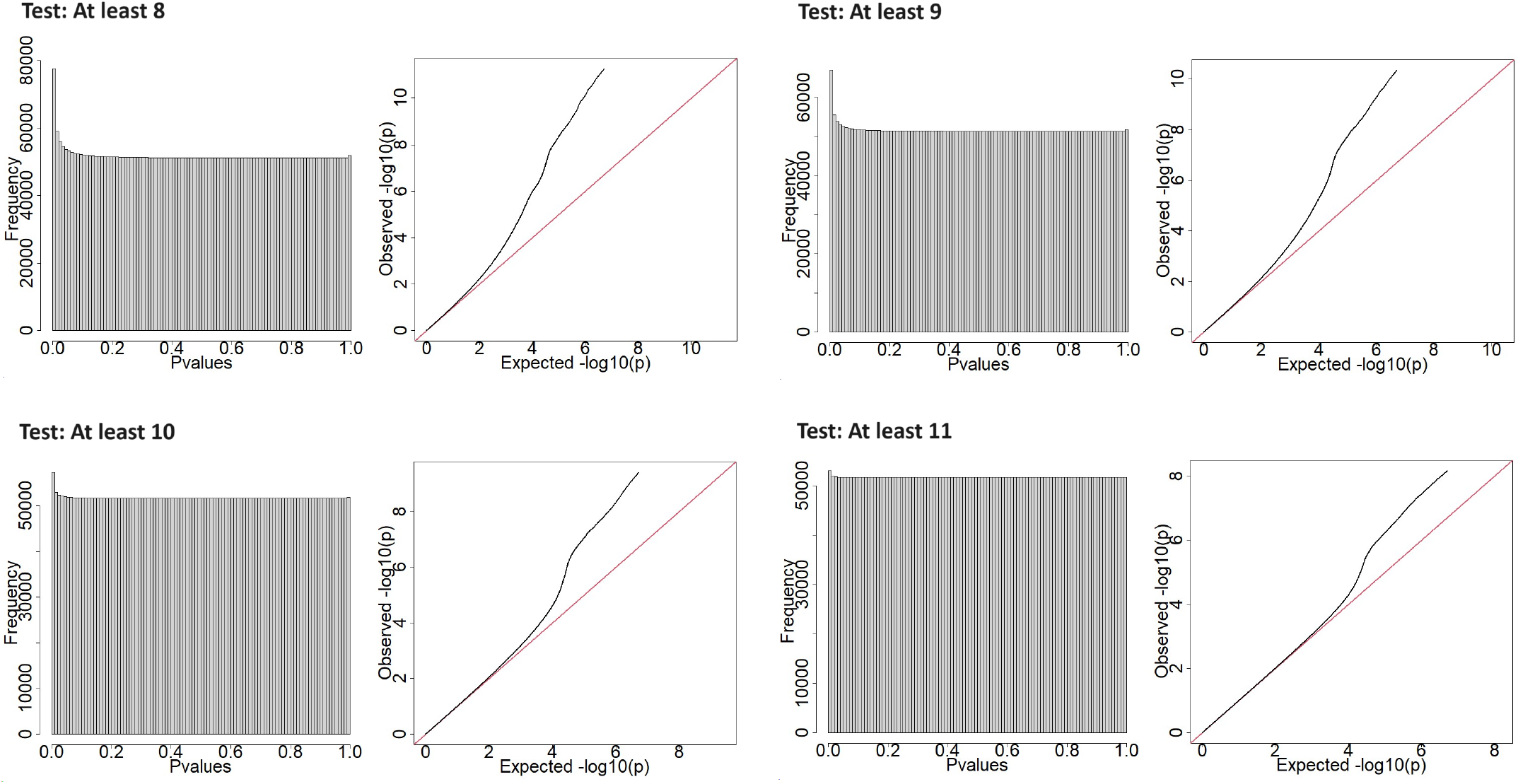
Quality control of the p-values distribution corresponding to the different composite hypothesis tests applied on the 14 psychiatric disorders GWAS. **Left:** Histogram of the qch_copula p-values. **Right:** Quantile-Quantile plot of the qch_copula p-values.

**Fig. 17.**
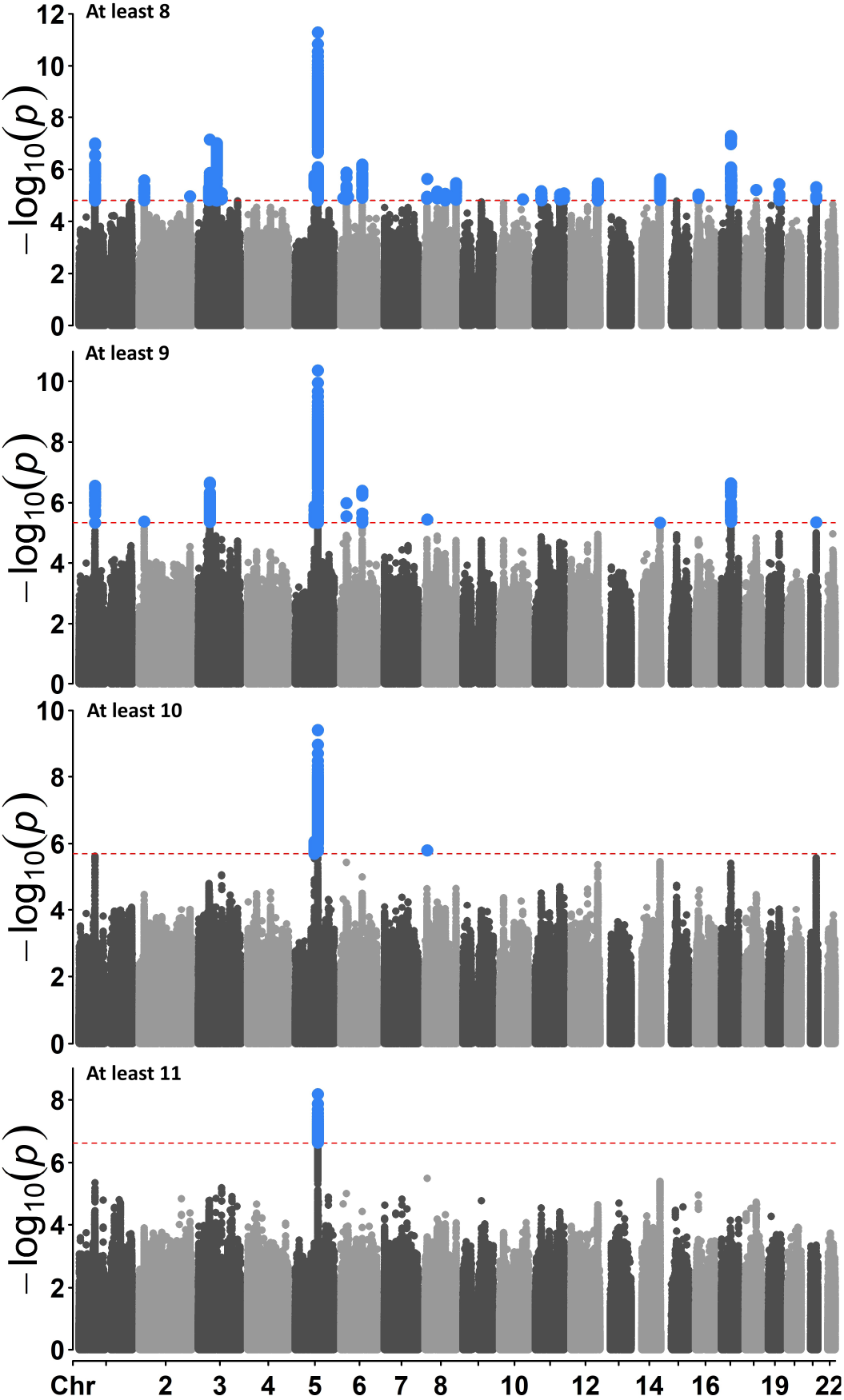
-log10(p-values) of the different composite hypothesis tests along the chromosomes applied on the 14 psychiatric disorders GWAS. From top to bottom, the composite hypothesis test identifies the SNPs associated with at least 8, 9, 10 and 11 disorders, respectively. The red dotted line represents the significance threshold at a nominal false discovery rate of 0.05. Significant SNPs are represented in blue.

**Fig. 18.**
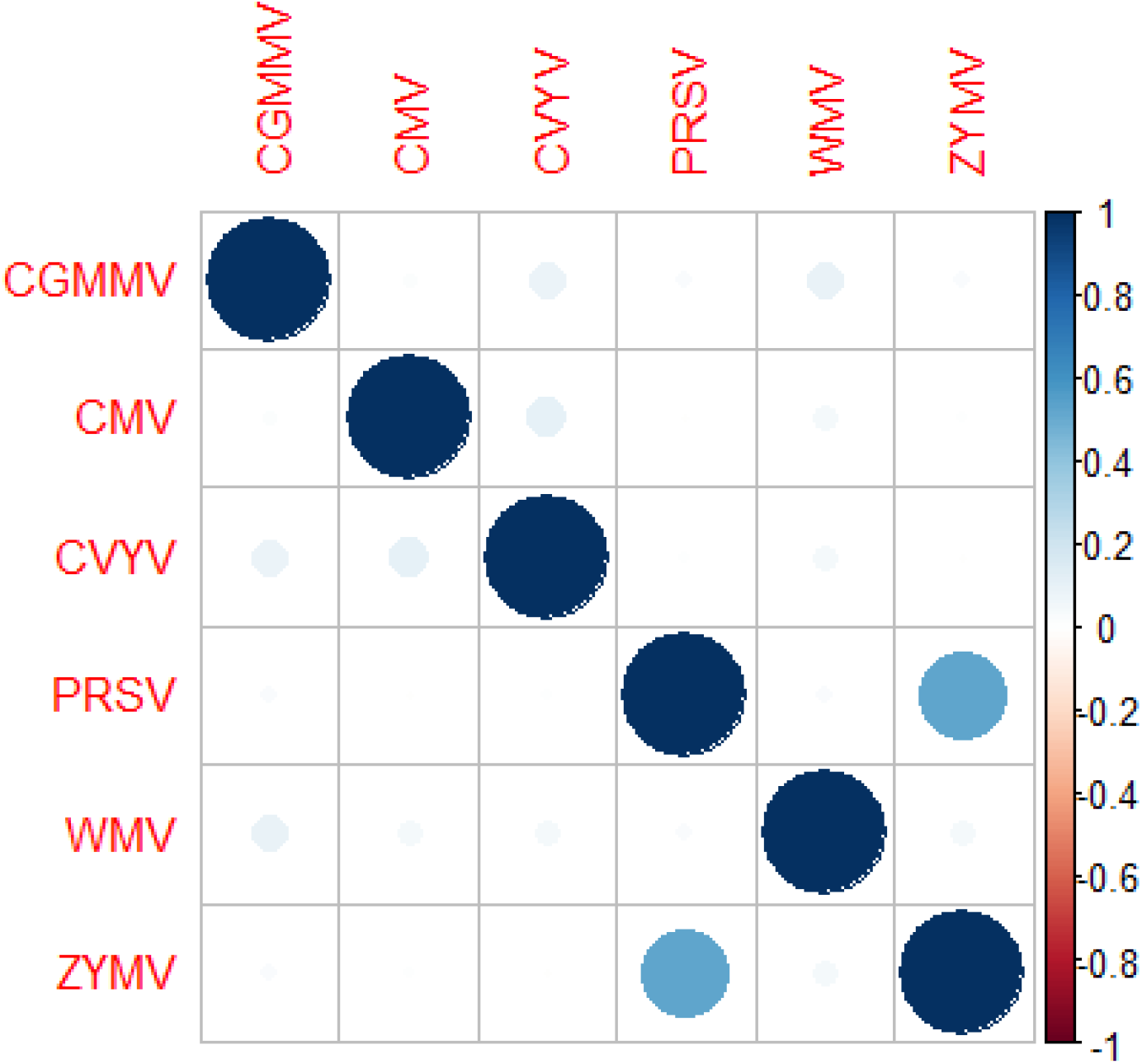
Estimated correlation matrix between the six GWAS on virus resistance.

**Fig. 19.**
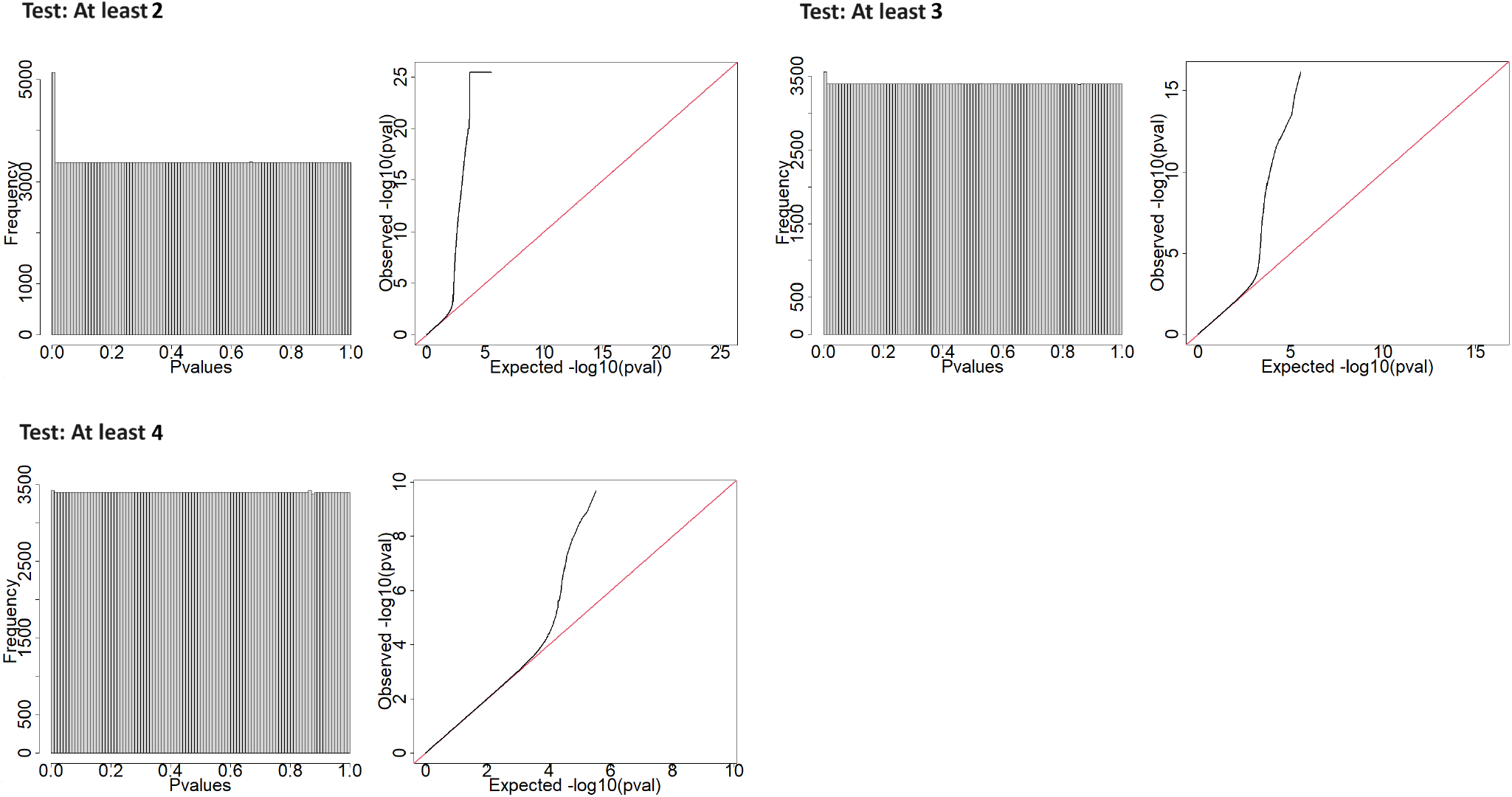
Quality control of the p-values distribution corresponding to the different composite hypothesis tests applied on the six viruses resistance GWAS. **Left:** Histogram of the qch_copula p-values. **Right:** Quantile-Quantile plot of the qch_copula p-values.

**Fig. 20.**
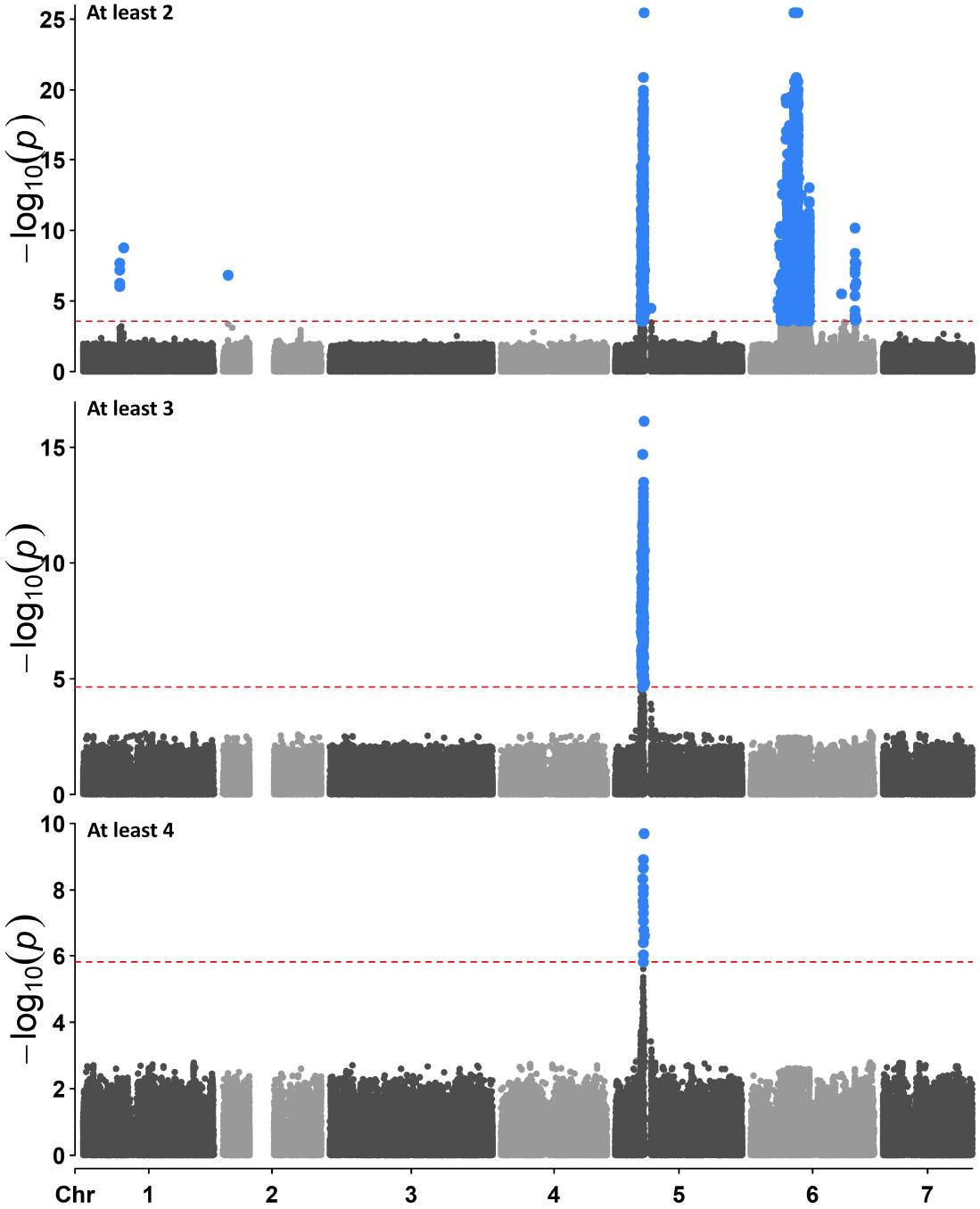
-log10(p-values) of the different composite hypothesis tests along the chromosomes applied on the six viruses resistance GWAS. From top to bottom, the composite hypothesis test identifies the SNPs associated with the resistance to at least 2, 3 and 4 viruses, respectively. The red dotted line represents the significance threshold at a nominal false discovery rate of 0.05. Significant SNPs are represented in blue.

## Footnotes

1 as available from the PGC website: https://pgc.unc.edu/for-researchers/download-results/

2 available from the following website: https://drive.google.com/drive/folders/1uFBeUwE-0UA0YmLTVQ9Ym_oU2HkHPdhk

